# The pioneer transcription factor ELF2 remodels the nucleosome near transcription start sites

**DOI:** 10.1101/2024.05.24.595749

**Authors:** Tianyi Xiao, Caillan Crowe-McAuliffe, Moritz Ochmann, Yepeng Du, Tianyun Hou, Yinan Chen, Fangjie Zhu, Fatemeh Seyednasrollah, Maria Osmala, Sakari Vanharanta, Ekaterina Morgunova, Patrick Cramer, Connor Rogerson, Christian Dienemann, Jussi Taipale

## Abstract

Pioneer transcription factors are DNA-binding proteins that can bind to nucleosomes in closed chromatin regions, exposing enhancers and promoters of genes for transcription. The action of these factors underpin stem cell pluripotency, cell reprogramming and differentiation. ELF2 is an ETS family pioneer factor that has a strong preference for oriented binding on nucleosomes at a composite head-to-tail dimeric sequence motif with a 2 bp spacing between the GGAA core elements. In this study, we investigated the interaction between ELF2 and a nucleosome using single-particle cryo-electron microscopy (cryo-EM). The ELF2-nucleosome structure shows two ELF2 proteins bound to the nucleosome cooperatively at superhelical location +4. The recognition ɑ4 helices of both ELF2 monomers dock into the major groove at the core GGAA motifs and unwrap approximately four helical turns of the DNA from the surface of the nucleosome. The unwrapping almost completely exposes one histone H2A:H2B dimer, which dissociates in a subset of particles to form an ELF2-bound hexasome. ChIP-seq combined with genome-wide motif mapping indicates that all unmethylated high-affinity ELF2 double motifs are occupied, indicating that ELF2 is able to use its binding energy to function as a pioneer factor both *in vitro* and *in vivo*. ELF2 binding is highly enriched downstream of transcription start sites (TSS) of highly expressed genes, with the motifs oriented in such a way that the TSS becomes accessible upon ELF2 binding. This mechanism is essential for cells, as mutation of ELF2 motifs result in decreased cell proliferation. Taken together, our results indicate that ELF2 can use its binding energy to open chromatin in an oriented fashion, facilitating nucleosome positioning and transcription initiation at transcription start sites.

## Introduction

A nucleosome is the basic unit of DNA packaging, it consists of ∼ 147 bp of DNA, and an octamer of two copies of each of histones H2A, H2B, H3 and H4. For a canonical nucleosome, DNA wraps 1 ¾ turns around the octamer in a left-handed manner (*1*). The formation of a nucleosome distorts the canonical B-form of DNA by bending and twisting the DNA towards the histone octamer in a superhelical trajectory. Assembly of nucleosomes on free DNA to form chromatin prevents transcriptional initiation (*2*), and generally decreases the accessibility of DNA for protein binding (*3*).

Transcription factors (TFs) are proteins that can recognise and bind to specific DNA sequences and regulate transcription (*4*). Whereas all TFs can bind free DNA, only a subset of TFs, the pioneer factors, can bind to nucleosomal DNA and activate transcription in closed chromatin regions (*5–8*). Pioneer factors are important in multiple cellular functions including development, differentiation and cellular reprogramming, and some pioneer factors have also been linked to pathological conditions such as cancer (*9–13*).

Numerous studies have investigated the structural basis of the interaction between TFs and DNA using X-ray crystallography, providing insights into the origins of specificity and affinity of protein-DNA interactions. However, despite the importance of the nucleosome and TFs in biology and medicine, much fewer structural studies have focused on the interactions between TFs and nucleosomes. Biochemical studies have established that there are five major modes of TF-nucleosome interaction, namely, a gyre spanning, periodic binding, dyad binding (*8*) and end binding modes (*8*, *14*, *15*), as well as an oriented binding mode, where the TF binding motif shows orientational preference relative to the nucleosome (*8*, *16*). Recently, several investigators, including us, have used cryo-EM to study pioneer factor-nucleosome interactions (*17–23*), identifying the molecular basis of the end-binding and periodic binding modes. However, the molecular basis of the oriented binding of TFs to nucleosomes has not been investigated previously. In this work, we have studied the mechanism by which the ETS family TF, ELF2, binds to nucleosomes in an oriented manner.

## Results

### ELF2 preferentially binds an oriented double motif at SHL+4

To investigate how ELF2 binds to DNA wrapped around a nucleosome, we first sought to determine the optimal binding motif and position of ELF2 on nucleosomes. We analyzed data derived from nucleosome consecutive affinity purification SELEX (NCAP-SELEX, Zhu et al., 2018) using MOODS (*24*) where the position of ELF2 motifs are searched for in NCAP-SELEX data. The motif search showed that ELF2 motifs are preferentially found at around 108 bp of the plus strand of the nucleosomal DNA (**Fig. 1a**). The majority of identified motifs constituted a double ETS motifs (CC**<u>GGAA</u>**GC**<u>GGAA</u>**GT) where the two GGAA core sequences are located in a head-to-tail orientation with a 2 bp gap, (referred to as ELF2-HT2 hereafter). The position of the ELF2-HT2 motif in the nucleosomal DNA coincides with superhelical location +4 (SHL +4) of the nucleosome (**Fig. 1b,1c**). This suggests that ELF2 preferentially binds to the nucleosome by recognizing an oriented double motif at SHL+4. This preference for a double motif at SHL+4 can also be observed across different classes (*25*) of ETS proteins such as class I (ERG, FLI1, FEV), class II (ELF1, ELF4, ELF5) and class III (SPIB) ETS factors (**Fig. S1**).

**Figure 1:**
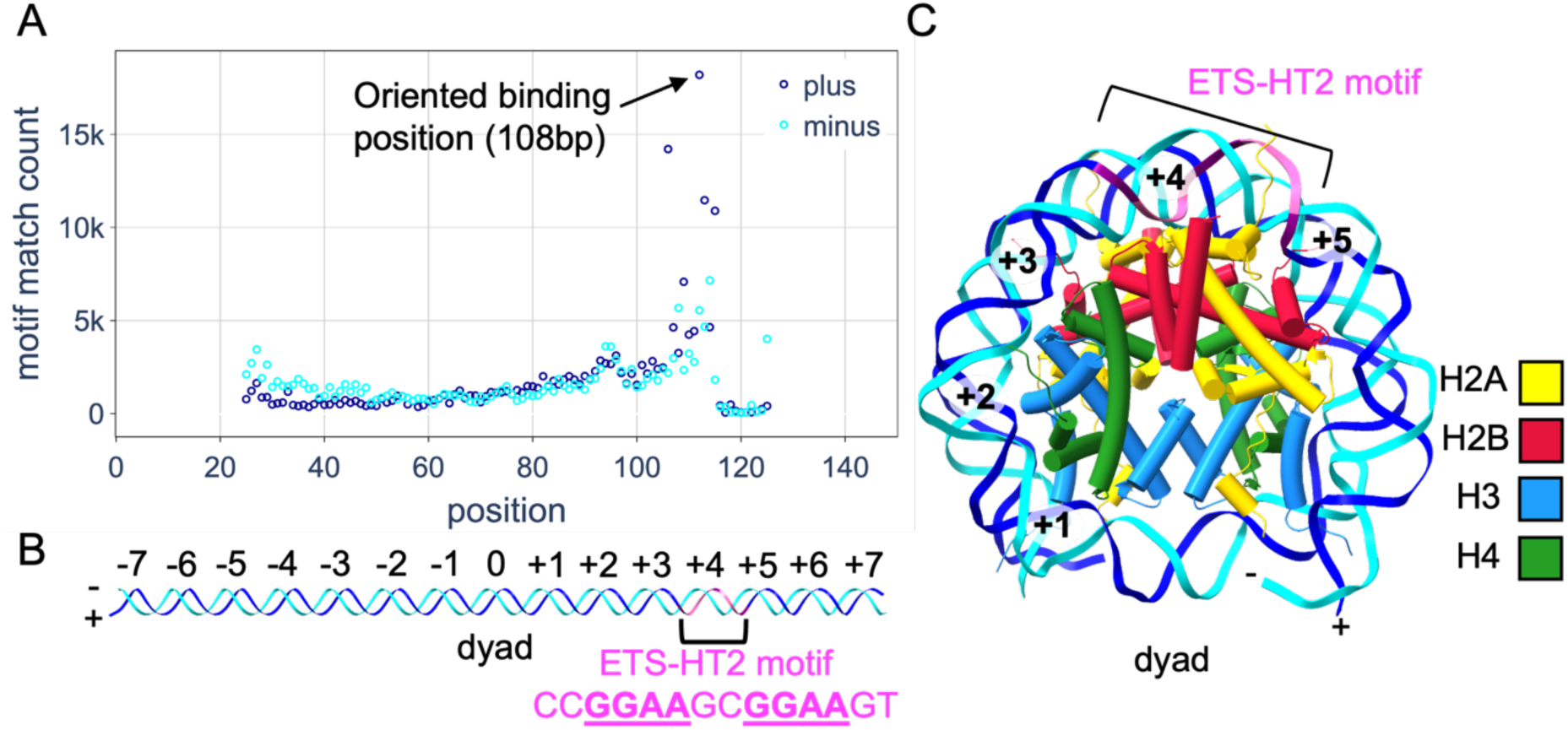
ELF2 displays high orientational and positional specificity on nucleosomal DNA. A: Preferred ELF2 binding position on the nucleosome from NCAP-SELEX data (Zhu et al., 2018). Dark and light blue circles represent binding in plus and minus strands, respectively. The oriented binding position of ELF2 is highlighted. B: DNA ligand design. Cartoon shows Widom 601 core sequence (147bp) with ETS-HT2 motif (CC**<u>GGAA</u>**GC**<u>GGAA</u>**GT) inserted in the most preferred binding position. The ETS-HT2 motif is shown in purple. Numbers indicate superhelical location. C: Positions of the two GGAA core binding sequences of ETS-HT2 on the nucleosome with superhelical locations labelled. The DNA sequence from 6FQ5 was mutated to the designed sequence using coot. The image is acquired using ChimeraX. Blue: H3; Green: H4; Yellow: H2A; Red: H2B.

In summary, our analysis of NCAP-SELEX data revealed that ELF2 prefers to bind to ELF2-HT2 motifs in the context of a nucleosome. Additionally, this binding preference appears to be a feature that can also be found for other ETS family members. This suggests that this binding mode may be a universal feature of ETS transcription factors in the context of a nucleosome.

### Cryo-EM structure of ELF2 bound to a nucleosome

We next aimed to elucidate structurally how ELF2 binds to the ELF2-HT2 motif at SHL+4. For this, we first reconstituted nucleosomes using a modified Widom 601 sequence (*26*) that contains the ELF2-HT2 motif at the preferred location (around SHL+4) as determined by the SELEX experiments (**Methods**, **Fig 1b, 1c**). An excess of purified ELF2 DNA binding domain (DBD) was then added and the sample was analysed with single-particle cryo-EM (**Methods**, **Fig. S2**, **Table S1**).

Initial cleanup of the cryo-EM data yielded a set of 1,179,230 nucleosome-like particles that were further analyzed (**Methods)**. After several rounds of 3D classification (**Fig. S2**), we obtained a reconstruction of a nucleosome at 3.8 Å global resolution (**Fig. S3a**) in which the density around SHL +4 substantially differs from the canonical nucleosome structure (**Fig. 2a**).

**Figure 2:**
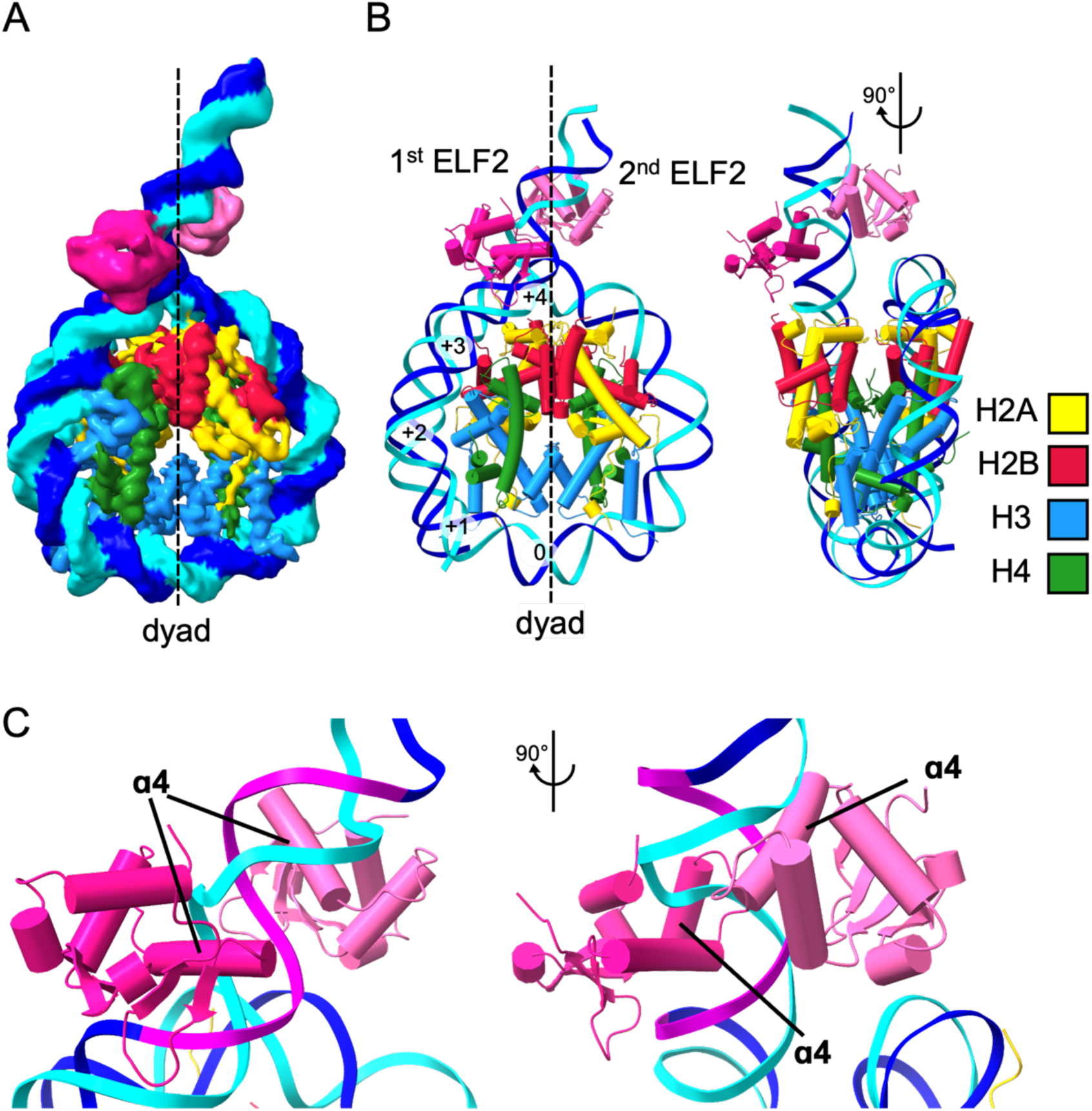
The structure of ELF2-bound nucleosome. A: ELF2-nucleosome composite cryo-EM map with dyad axis shown. The map shown is a composite map from a global refinement and local refinement on ELF2 (**Fig. S2**). B: Front and side view of ELF2-nucleosome model. C: Local ELF2-nucleosome interaction. ɑ4 helices of ELF2 DBDs are docked into the CC**<u>GGAA</u>**GC**<u>GGAA</u>**GT binding motif (purple). Right image is 90° rotation around the y axis from the left.

First, a long rod-like density pointing away from the nucleosome could be explained by fitting approximately B-form DNA that exits the nucleosome at SHL+4 (**Fig. 2b**). Additionally, two extra densities bound to this DNA were present. Due to flexibility in this region, local refinement was used to improve the resolution (**Methods**). This resulted in a 5.4 Å reconstruction of this region (**Fig. S3b**, **Fig. S4a**), which was sufficient to fit two copies of the ELF2 DBD. Fitting of the DBDs placed the recognition helices of both ELF2 DBDs in the major grooves of the nucleosomal DNA at the expected GGAA sites, with the two proteins bound to DNA on opposite sides of the DNA helix (**Fig. 2c**). Of the initial particle set, only ∼ 4% represented a free nucleosome without ELF2 bound (**Fig. S2**), indicating that under the conditions tested, ELF2 has a high occupancy, and its binding to nucleosomes is relatively stable.

On a fully wound nucleosome, the first ELF2 binding site (the one that is closer to the nucleosome in the structure) is largely accessible from the solvent side, whereas the second site is occluded in such a way that an ELF2 bound to it would clash with the other DNA gyre (**Fig. S5**), suggesting a consecutive binding mechanism for the two ELF2 proteins. The structure we obtained also suggests a mechanism for the orientational preference of ELF2 nucleosome binding. The ELF2 monomer bound to the second site is oriented in such a way that a lysine, K229, is located near the phosphate backbone of the other gyre of DNA at SHL - 4 (**Fig. 3a**). The side-chain of this lysine is not resolved in the structure, but it could reach within 4.5 Å of the phosphate at base 35, indicating a potential electrostatic interaction.

**Figure 3:**
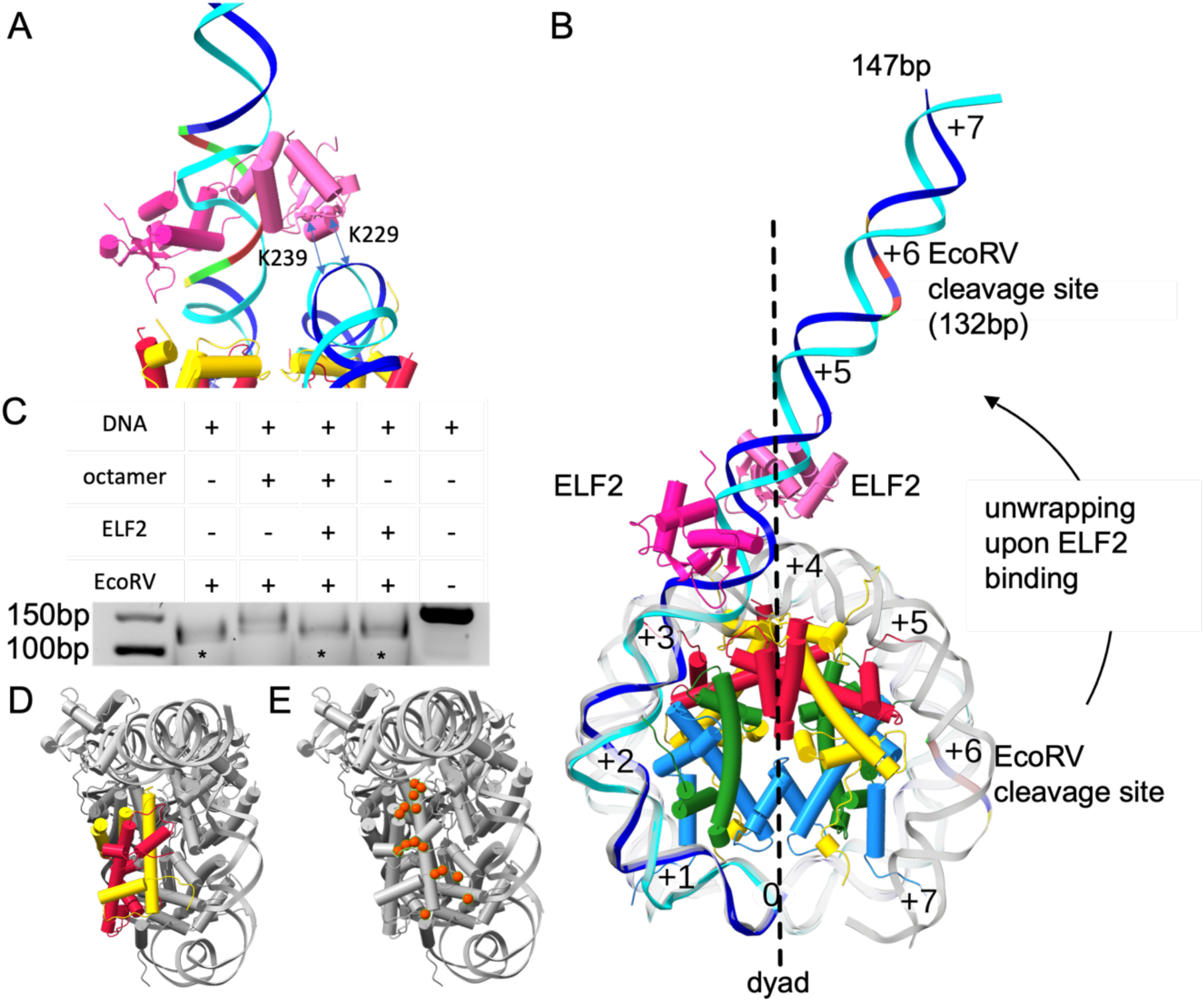
ELF2 unwraps nucleosome, resulting in hexasome formation. A: Side view of ELF2-nucleosome interaction with K229 and K239 highlighted with the potential phosphate backbone interaction indicated by arrowed lines. B: ELF2-nucleosome model overlaid on a canonical nucleosome (6FQ5 DNA shown in gray). Dyad axis is labelled. The superhelical locations on nucleosomes are labelled. The engineered EcoRV cleavage site is indicated. The model was extended in Coot with ideal B-form DNA on the unwrapped end to represent the full 147 bp DNA ligand. C: Restriction enzyme digestion assay. Asterisks indicate lanes with faster-migrating DNA bands (DNA cleaved by EcoRV). Samples were denatured before gel electrophoresis. D: Top view of ELF2-nucleosome model, exposed H2A:H2B dimer exposed are highlighted. E: Top view of ELF2-nucleosome model showing contacts between H2A:H2B and DNA that are lost after unwrapping, indicated by orange spheres.

### ELF2-binding unwraps the nucleosome

The long DNA density exiting the nucleosome at SHL+4 indicates that a large portion of nucleosomal DNA has detached from the octamer in our structure. Indeed, there is no density for wrapped nucleosomal DNA continuously extending from SHL+4 to +7 (**Fig. 2a**) showing that approximately 40 bp of DNA have been fully detached from the nucleosome (**Fig. 3b**). Two of the helical turns (at SHL +4 and +5) of the unwrapped DNA are still resolved in the structure, whereas the rest remains unresolved due to flexibility (**Fig. S4b**). Weak and non-continuous residual density for nucleosomal DNA from SHL +5 to +7, probably originating from a strand of free DNA that binds the exposed H2A:H2B dimer *in trans*, could still be observed at low display thresholds (**Fig S4c**). In addition, hexasomes are observed in a small population of particles (**Fig. S4d**), where one H2A:H2B dimer is lost where the DNA is detached. Altogether, the structure suggests that binding of ELF2 to SHL+4 of the nucleosome induces unwrapping to the nucleosomal DNA from SHL +4 to +7.

To confirm this, we tested unwrapping of nucleosomal DNA by ELF2 binding *in vitro*. For this, we introduced an EcoRV cleavage site at around SHL+6 of the nucleosomal DNA (**Fig. 3b**) and tested the ability of EcoRV to cleave the DNA in the context of a nucleosome and in the presence or absence of ELF2. EcoRV was chosen as it requires >300° of access around the DNA helix to interact (*27*) and thus only recognizes and cleaves DNA that is fully detached from the nucleosome. In the context of a nucleosome and without ELF2, the nucleosomal DNA is protected from EcoRV digestion at SHL+6 as expected for nucleosomal DNA that is stably wrapped around the histone octamer (**Fig. 3c**). However, EcoRV efficiently cleaves nucleosomal DNA when the nucleosome was incubated with ELF2 before adding the nuclease (**Fig. 3c**), consistent with ELF2-induced unwrapping of DNA.

Altogether, these experiments show that binding of ELF2 to an ELF2-HT2 motif at SHL+4 of a nucleosome leads to unwrapping of nucleosomal DNA from SHL+4 to +7. The extend of DNA unwrapping observed upon ELF2 binding exceeds the unwrapping reported for pioneer transcription factors such as SOX2, CLOCK-BMAL1 and OCT4 (*17*, *20*, *21*, *23*). The data further suggest that ELF2 binding to ELF2-HT2 motifs located in a nucleosome may lead to chromatin opening. Thus, ELF2 may act as a pioneer transcription factor when bound to an ELF2-HT2 motif.

### ELF2 binding can release an H2A:H2B dimer

Unwrapping of nucleosomal DNA due to binding of ELF2 also exposes one of the H2A:H2B dimers to solvent (**Fig. 3d**). Almost all contacts between this H2A:H2B dimer and DNA are lost compared to those observed in the canonical nucleosome structure (**Table S2**; **Fig. 3e**). Upon ELF2 binding, the predicted binding surface between the H2A:H2B at SHL+5 and the nucleosome decreased from 1877.2 Å^2^ to 279.6 Å^2^. The solvation free energy of this interaction decreased from -20.8 kcal/mol to -5 kcal/mol. This suggests that the binding of this H2A:H2B dimer to the nucleosome is weakened upon DNA unwrapping by ELF2. Consistently, we observed a class of 23,778 particles that was structurally similar to the major class except that it was missing the H2A:H2B dimer at the site of ELF2-induced DNA unwrapping (**Fig. S4d**). Thus, unwrapping of nucleosomal DNA by ELF2 binding can lead to release of an H2A:H2B dimer and formation of a hexasome.

### ELF2 binds downstream of the TSS

Pioneer transcription factors modulate transcription by binding to enhancers or gene promoters (*28*). We therefore investigated the localization of ELF2-HT2 motifs in the genome and whether genomic regions containing ELF2-HT2 motifs are bound by ELF2 *in vivo*. We first determined where ELF2 can bind to the genome and searched for ELF2 and ELF2-HT2 motifs in the human genome. We found that ELF2 single motifs are enriched in gene promoters and enhancers, while ELF2-HT2 motifs are enriched in gene promoters (**Fig. S6a,b**). In particular, we found an enrichment for ELF2-HT2 motifs at 5-10 bp downstream from TSS (**Fig. 4a**). The enrichment was specific for the minus-strand of the respective genes, which shows that ELF2- HT2 motifs downstream of the TSS occur in a specific orientation relative to the TSS and the direction of transcription. Single ELF2 motifs show weaker orientational preference relative to the TSS (**Fig. 4a**). The extent of oriented enrichment of ELF2-HT2 motifs downstream of the TSS is similar to YY1, a TF known to have strong oriented binding preference 20-30 bp downstream of the TSS (*29*, *30*).

**Figure 4:**
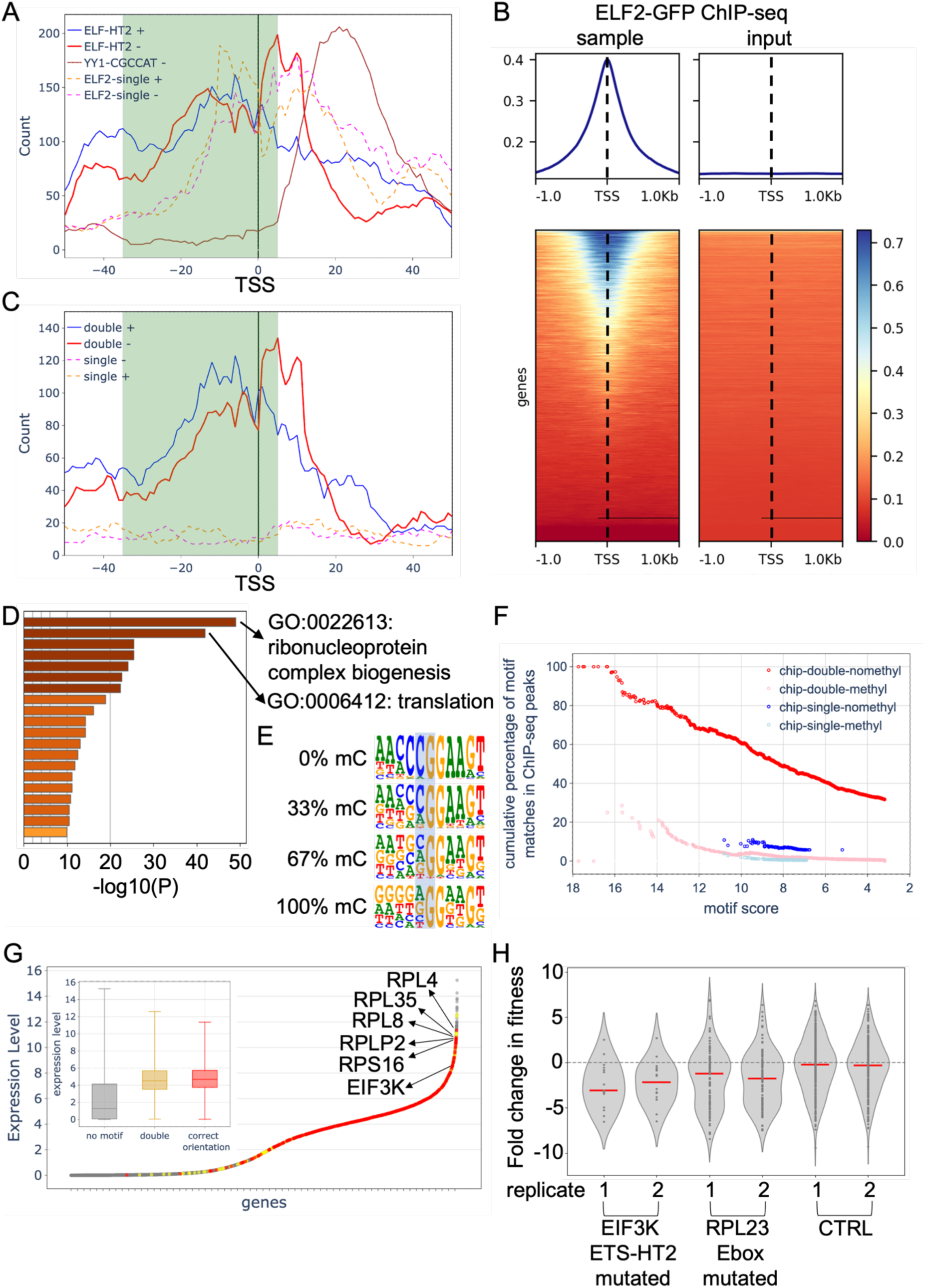
Potential role of ELF2-HT2 motif in facilitating transcription initiation. **A**: Enrichment of ETS-HT2 motifs near TSS. Metagene plot of the enrichment of the indicated TF motifs relative to the TSS positions in the human genome (hg38). Distance from TSS against motif count in a 10 bp sliding window is shown. Enrichment for plus and minus strand are shown separately. DNA that would be made accessible by nucleosome unwrapping is highlighted by green shading. **B**: heatmap of ChIP-seq peaks relative to TSS plotted using deeptools. TSS plotted in this plot are all TSS in hg38. Dotted lines show the TSS position. **C**: positioning of ETS-HT2 motifs (ACCC**<u>GGAA</u>**GC**<u>GGAA</u>**GT) found from ChIP-seq relative to TSS in human genome hg38. The motif count is shown in a 10bp sliding window. The double motif is enriched downstream from TSS pointing towards TSS. Green shaded region shows the DNA that would be made accessible from nucleosome by ELF2 binding. single ELF2 motifs found from ChIP-seq are shown as dotted lines. **D**: Gene ontology analysis of ETS-HT2 motifs discovered from ChIP-seq. **E**: Motif logo of ELF2 binding sequence in mC-SELEX under different methylcytosine (mC) levels. **F**: Scatter plot of cumulative percentage of motif matches in ChIP-seq peaks against MOODS motif score. For the double motifs, red shows neither CG sites are methylated and pink shows both CG sites are methylated. **G**: Scatter plot of gene expression levels in K562 grouped by not having a double ETS motif at TSS (grey), having double motifs near TSS but more than 20bp away (yellow) and having ETS double motif within 20bp of TSS and in the orientation that the binding would make TSS accessible (red). Top 5 genes with ETS-HT2 double motif within 20 bp of TSS are labelled. Expression level of EIF3K is labelled. Box plot shows gene expression levels in K562, with same colouring scheme as the scatter plot. The box shows upper quartile and lower quartile, the line within the box is the median, whiskers show the minimum and maximum. The ETS-HT2 double motifs are found from ChIP-seq data. **H**: violin plot of cell fitness assay. negative control, positive control (E-box knockout) and EIF3K ELF motif knockout is shown.

To confirm that ELF2 also binds downstream of the TSS at gene promoters, we generated two clonal K562 cell lines where a GFP-tag is knocked into the C-terminus of ELF2 (see **Methods**) and then performed ChIP-seq using a GFP-antibody. As expected from the motif enrichment at promoters, the ChIP signal was mostly located at gene promoters (**Fig. S6c**) where it peaks centered on the TSS (**Fig. 4b**). Motif discovery using BPNet (*31*), a deep learning model that can generate motifs based on contribution scores, resulted in identification of both a single ELF2 motif and the ELF2-HT2 motif as the most enriched double ETS motif in the peaks (**Fig. S7**). When searching for ELF2-HT2 motifs in the ChIP-seq peaks, we find oriented ELF2-HT2 double motifs with similar enrichment compared to the genome-wide motif search (**Fig. 4c**). This confirms that genomic regions downstream of the TSS that contain ELF2-HT2 motifs are also bound by ELF2 *in vivo*.

Altogether, we show that ELF2-HT2 motifs are enriched downstream of the TSS and that these regions are also bound by ELF2 *in vivo*. This suggests that ELF2 binding to ELF2-HT2 motifs located at gene promoters ELF2 may cause unwrapping of promoter DNA that is covered by nucleosomes and therefore inaccessible for transcription initiation. Consistent with this, ELF2 was found to generate nucleosome free regions of ∼150 bp around its binding sites *in vivo* (*8*).

### ELF2 pioneer activity is associated with high expression levels

Accessibility of the promoter region for binding of the Pol II PIC is a prerequisite for efficient transcription initiation and the presence of a nucleosome that overlaps with the promoter has been shown to be detrimental to gene expression (*32*, *33*). We therefore checked if ELF2 binding to gene promoters is associated with higher levels of transcription. A gene ontology analysis of genes with ELF2 ChIP-seq peaks downstream of their TSS showed an enrichment for the GO terms “translation” and “ribonucleoprotein complex biogenesis”, which usually describe highly expressed genes (**Fig. 4d**).

To assess whether ELF2 also acts as a pioneer factor *in vivo*, we determined the fraction of ELF2 double motif matches that are occupied (located within ChIP-seq peaks). As CpG methylation of the CG dinucleotide in the ETS consensus sequence CGGAA is known to inhibit DNA binding of ETS factors (*34*, *35*), **Fig. 4e**), we considered doubly methylated and fully unmethylated motif matches separately by combining the motif match positions with K562 cell whole-genome bisulfite sequencing data from GEO (*36*). This analysis revealed that all unmethylated high-affinity ELF2 double motifs are occupied, and that the occupancy decreased as a function of motif match score (**Fig. 4f**). By contrast, methylated double motif matches, or single motif matches were occupied much more infrequently, consistent with a central role for the binding energy of TFs for pioneer activity (*37*).

We next determined whether the presence of oriented ELF2-HT2 motifs at the promoter is associated with high levels of gene expression using DEPMAP gene expression data from K562 cells (*38*). We first identified all genes containing an ELF2 ChIP-seq peak and an ELF2-HT2 double motif within 20 bp of their TSS. Almost all such genes are expressed with expression levels significantly (p = 4.62 x 10^-126^, Student’s T-test) higher than genes that do not contain ELF2 ChIP-seq peaks or an ELF2-HT2 motif within 20 bp of TSS (**Fig. 4g**). Interestingly, the top five expressed genes with ELF2-HT2 motifs are all ribosomal protein genes (**Fig. 4g**). This suggests that an oriented ELF2-HT2 motif and ELF2 binding at gene promoters is associated with high gene expression levels.

To confirm that oriented ELF2-HT2 motifs are important for maintaining gene expression levels, we chose a highly expressed (**Fig. 4g**) and essential target gene, EIF3K, that contains two high-affinity ELF2-HT2 motif matches, and mutated the double motifs using a competitive precision genome editing assay (CGE, (*39*)). We then measured the fitness of these cells by comparing the number of each flanking sequence tag between day 2 and day 8. Mutation of the ELF2-HT2 motifs downstream of the TSS of the EIF3K gene resulted in decreased cell fitness (**Fig. 4h**), confirming that ELF2-HT2 motifs and therefore ELF2 binding at this promoter contributes to gene expression.

Altogether, the presence of ELF2-HT2 motifs downstream of the TSS of a gene promoter is highly correlated with high expression levels. Further, these data show that the integrity of the binding sequence is crucial for maintaining expression of genes containing an ELF2-HT2 motif at their promoter. Thus, we propose that ELF2 binding to ELF2-HT2 motifs at gene promoters supports efficient transcription, likely by unwrapping promoter DNA from nucleosomes that is otherwise masked.

## Discussion

In this work, we have solved the structure of ELF2 bound to the nucleosome. We found that by binding in a head-to-tail dimeric fashion, ELF2 protein unwraps the nucleosome by four superhelical turns, exposing 40 bp of free DNA and the H2A:H2B interface. Structures of many pioneer factor-nucleosome interactions have been described in recent publications (*17–23*). Most pioneer factors described bind near to the end of nucleosomal DNA. However, in this study, we examined ELF2, which prefers to bind at around SHL +4, opposite of the dyad. Compared to some other pioneer factors such as CLOCK (*20*), ELF2 binds to DNA further into the nucleosome, and unwraps more DNA. The ELF2 dimer motif that is preferentially bound on a nucleosome is enriched downstream of the TSS in such a way that binding to the site would expose the TSS, suggesting that the unwrapping mechanism we discovered may promote the initiation of transcription.

### Biological role of ETS factors

The ETS family is a large TF family found only in animals. Several ETS family members have central roles in development and disease. For example, the class I ETS factors ERG, ETV1 and ETV4 are linked to prostate cancer (*40*, *41*). The ELF family, in turn, has been reported to have important roles in immune and blood cell development and function (*42*). In addition, double motifs for ETS have been suggested to be important for transcription initiation (*43*). We show here that the nucleosome-bound ELF2-HT2 motif is bound by many members of this family (**Fig. S1**), suggesting that ELF2, and ETS factors more broadly would function as pioneer factors, destabilizing nucleosomes and increasing DNA accessibility and transcription.

### Nucleosome unwrapping & chromatin modifications

The mechanism of nucleosome unwrapping observed in this study is based on binding energy, as it involves no enzymes or ATP hydrolysis. Consistently, we found that also in cells, all unmethylated high-affinity double ELF2 motif matches were occupied, indicating that nucleosomes are not able to occlude these sites *in vivo*. By contrast, unmethylated maximal affinity single ELF2 motifs were occupied only ∼10% of the time, consistent with prior observations for many individual TF motif matches (*44*, *45*). Correlative evidence from DNase I footprinting indicates that many typical open chromatin regions contain five to six highly occupied TF motif positions (*46*). The particular TF binding events that cause open chromatin formation in cells still remain largely unknown. However, our results suggest that even a single locally cooperative TF binding event (*47*, *48*), driven by mass action, may be sufficient for complete or partial displacement of nucleosomes.

As the two ELF2 proteins bind to opposite sides of DNA, they cannot simultaneously bind to nucleosomal DNA. We propose that one ELF2 binds to the first binding site, pulling DNA away from the histones without completely releasing the DNA. This allows the second ELF2 to bind to the second site, leading to unwrapping of the DNA end. After ELF2 binding and nucleosome unwrapping, DNA stays in an unwrapped state because the second ELF2 prevents re-wrapping of the nucleosome. This mechanism would explain how ELF2 can access an optimal motif present at an optimal position. However, in the genome, nucleosome positions vary, and the first binding site will often be occluded. Even in that case, the binding of the ELF2 proteins to opposite sides of DNA would still ensure that one of the sites would be accessible for binding. We propose that such motifs, which enable binding to both sides of DNA, will be a general mechanism that will allow pioneer factors to access nucleosomal DNA irrespective of the rotational positions of their recognition sequence(s). In addition to direct binding to wrapped DNA, ELF2 can also bind to nucleosomes affected by spontaneous local conformational fluctuations (DNA breathing) that temporarily expose nucleosomal DNA ends (*15*, *49*, *50*).

### Oriented binding

Wrapping of DNA to nucleosomes breaks the rotational (pseudo)symmetry of the DNA molecule, allowing some TFs, such as ELF2, to prefer to bind to nucleosomes in an oriented fashion. Prior to the present work, the mechanism of oriented binding was not understood, as it has not been studied in structural detail. We find here that ELF2 has a lysine residue that can reach out to the phosphate backbone of the opposite DNA gyre. It is therefore speculated that interacting with the phosphates stabilizes ELF2 binding on the second binding site, promoting ELF2 dimer binding in this preferred orientation. If the motif is found in reverse complement (TTCC instead of GGAA), the ELF2 protein would rotate 180 degrees, orienting the lysine away from the other DNA gyre. Oriented binding allows TFs to position nucleosomes precisely relative to their binding motif. We propose that this mechanism may be important in positioning of nucleosomes relative to the TSS, and possibly also in setting the boundaries of open chromatin regions in general.

### Hexasome and chromatin remodelling

We found here that after ELF2 binding unwraps a nucleosome, the octamer becomes unstable and in some cases dissociates partially, with one H2A:H2B dimer dissociating, which yields a hexasome. Hexasomes comprise around 110 bp DNA wrapped around six histones (*51*). It has been shown previously that hexasomes recruit chromatin remodelers such as INO80 (*52*, *53*). Our finding that ELF2 binding can lead to hexasome formation thus suggests a potential novel mechanism by which TFs can recruit chromatin remodelers and increase transcription rates.

Interestingly, AlphaFold3 cannot produce a high confidence structure with full octamer, DNA and two ELF2 proteins as an input. It instead predicts a physically unrealistic structure of a fully wrapped nucleosome, with the two ELF2 proteins bound to their sites in a highly strained conformation (**Fig. S8**). However, an input of hexasome, DNA and two ELF2 proteins yields a model that is similar to that obtained by Cryo-EM (**Fig. S8**). This result increases confidence in the existence of the ELF2-hexasome complex and suggests that computational modelling can in optimal cases yield realistic TF-nucleosome structures, even if these represent intermediates.

### Role of oriented binding in transcriptional regulation

TFs recognise specific motifs on DNA and regulate transcription via transcriptional activation domains (*54*). Most TFs do not require strict positioning in promoters or enhancers (*30*). For example, single ETS motifs with core sequence GGAA are broadly located upstream of the TATA box, between TATA box and TSS, and downstream of TSS. By contrast, we find here that the ETS-HT2 dimer motifs in ChIP-seq peaks are preferentially oriented and located downstream of the TSS positioned in such a way that the TSS DNA sequence would be released from a nucleosome (**Fig. 4a, 4c**).

Unwrapping of nucleosomes by ELF2-DNA interaction generates free DNA for other non-pioneer TF to bind to, and exposes histone H2A:H2B surfaces that are normally buried by nucleosomal DNA. Based on these findings, we propose a model where the ELF2 or other ETS factor binding at the ETS-HT2 sites positioned downstream of a TSS increases transcription by a novel mechanism consisting of three separate actions: 1) increasing DNA accessibility at the TSS, 2) positioning the +1 nucleosome, and 3) modifying the +1 nucleosome by formation of a hexasome (**Fig. 5**). The proposed novel mechanism differs from the classical activator model in two ways: it critically depends on precise positioning and orientation of the motif, and enables TFs without transcriptional activator domains to activate transcription.

**Figure 5:**
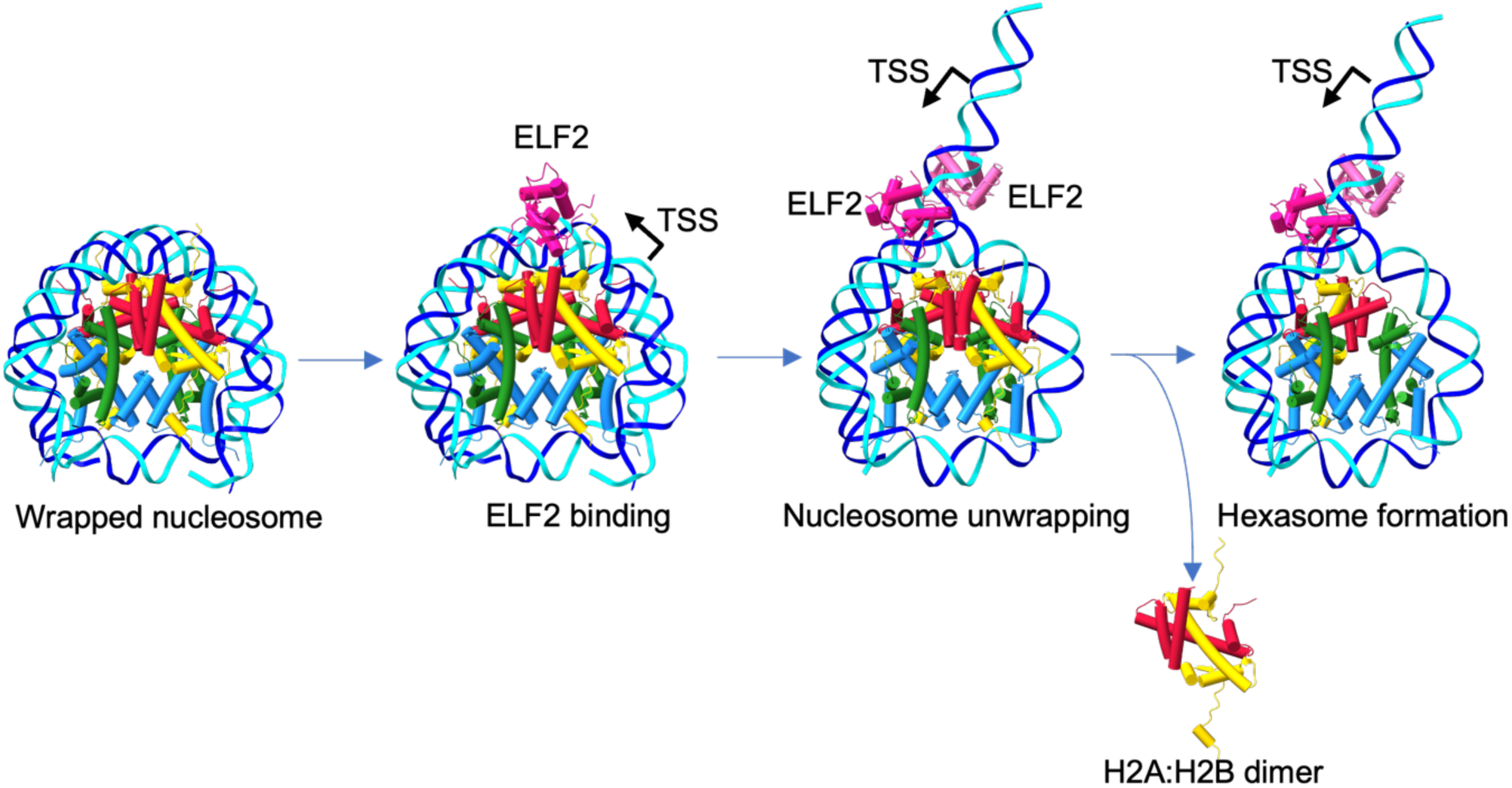
schematic of DNA unwrapped by ELF2. Model of DNA unwrapping and transcriptional regulation by ELF2. ELF2 binding unwraps the nucleosome by about 40bp. ELF2 motifs are enriched near the TSS and oriented so that unwrapping after binding would make the TSS accessible. In addition, ELF2 promotes formation of hexasomes.

In summary, we describe here the structure of an ETS factor ELF2 bound to nucleosomes as a dimer. Our work is the first structural study of a TF accessing nucleosomes in an oriented fashion, binding in a mode that enables precise control of nucleosome positioning and local DNA accessibility.

## Methods

### DNA ligand design

The nucleosomal DNA sequence used for the structural study included a dual ELF2 binding motif (CC<u>GGAA</u>GC<u>GGAA</u>GT) inserted into the preferred binding position within a Widom 601 core sequence (147 bp). The full sequence of the DNA (`DNA1`) used in the structural studies (ETS-HT2 motif <u>underlined</u>) was: 5’- CTGGAGAATCCCGGTCTGCAGGCCGCTCAATTGGTCGTAGACAGCTCTAGCACCGCTTAAAC GCACGTACGCGCTGTCCCCCGCGTTTTAACCGCCAAGGGGATTACT<u>CCGGAAGCGGAAGT</u>TG TATGTCAGATATATACATCCTGT-3’

The following three DNA sequences, each containing a single EcoRV site were also constructed for the restriction digestion analyses. The sequence (`DNA2`) with ELF2-HT2 motif (<u>underlined</u>) and EcoRV site (**bold**) was: 5’- CTGGAGAATCCCGGTCTGCAGGCCGCTCAATTGGTCGTAGACAGCTCTAGCACCGCTTAAAC GCACGTACGCGCTGTCCCCCGCGTTTTAACCGCCAAGGGGATTACT<u>CCGGAAGCGGAAGT</u>TG TATGTCA**GATATC**TACATCCTGT-3’

The sequence (DNA3) with only the first ELF2 motif (underlined) and EcoRV site (**bold**) was: 5’- CTGGAGAATCCCGGTCTGCAGGCCGCTCAATTGGTCGTAGACAGCTCTAGCACCGCTTAAAC GCACGTACGCGCTGTCCCCCGCGTTTTAACCGCCAAGGGGATTACTCCGGAAGCGTTAGTTG TATGTCA**GATATC**TACATCCTGT-3’ and the sequence (DNA4) with only the second ELF2 motif (underlined) and EcoRV site (**bold**) was: 5’- CTGGAGAATCCCGGTCTGCAGGCCGCTCAATTGGTCGTAGACAGCTCTAGCACCGCTTAAAC GCACGTACGCGCTGTCCCCCGCGTTTTAACCGCCAAGGGGATTACTCCATAAGCGGAAGTTG TATGTCA**GATATC**TACATCCTGT-3’

The fragments were constructed from one single-stranded full-length oligonucleotide using PCR (Phusion polymerase, 35 cycles, annealing 57°C 10s, elongation 72°C 36s, denaturation 98°C 15 s) with the following Forward primer (common) CTGGAGAATCCCGGTCTGC. The reverse primer Rev1 ACAGGATGTATATATCTG was used for DNA1, and Rev2 ACAGGATGTAGATATCTG for DNAs 2-4. All DNAs 1-4 were purified by anion exchange using ResourceQ 1 ml column (Cytiva), followed by isopropanol precipitation.

### ELF2 protein expression & purification

DNA-binding domain fragment of human ELF2 (encoding amino-acids 182-326; sequence: TQQSPISNGSPELGIKKKPREGKGNTTYLWEFLLDLLQDKNTCPRYIKWTQREKGIFKLVDSKAVSKLWGKHKNKPDMNYETMGRALRYYYQRGILAKVEGQRLVYQFKDMPKNIVVIDDDKSETCNEDLAGTTDEKSLE RVSLS) were cloned by Genscript and purified by Protein Science Facility at Karolinska Institutet.

### Histone octamer expression and purification

*Xenopus laevis* histones H2A, H2B, H3, and H4 were expressed and purified as described previously (*55*, *56*). In brief, each histone was expressed in BL21 (RIL) DE3 (Agilent) for 3 h at 37°C induced by 1 mM IPTG. Histones were extracted and purified from inclusion bodies. Cell pellets were resuspended (in 50 mM Tris-HCl pH 7.5 (at 20°C), 100 mM NaCl, 1 mM EDTA, 2 mM DTT, 0.284 µg ml^−1^ leupeptin, 1.37 µg ml^−1^ pepstatin A, 0.17 mg ml^−1^ PMSF, and 0.33 mg ml^−1^ benzamidine), lysed by sonication and pelleted by centrifugation. The pellet was washed repeatedly (with and without 1% (v/v) Triton X-100), before the histones were extracted from the inclusion bodies using DMSO followed by unfolding buffer (7 M guanidine hydrochloride, 20 mM Tris-HCl pH 7.5 (at 20°C), 10 mM DTT). The resuspended histones were buffer exchanged (using HiPrep 26/10 Desalting column (Cytivia) to 7 M deionized urea, 20 mM sodium acetate pH 5.2, 200 mM NaCl, 1 mM EDTA, 2 mM DTT) and purified with a HiTrap Q HP column (Cytivia) followed by a HiPrep SP FF 16/10 column (Cytivia). Fractions containing purified histones were dialyzed against 15 mM Tris-HCl pH 7.5 (at 20°C) and 2 mM DTT before being aliquoted, snap-frozen, and lyophilized before storage at -70°C. Histone octamer were formed as described (*55*). Histones H2A, H2B, H3, and H4 were resuspended in unfolding buffer (7 M guanidine hydrochloride, 20 mM Tris-HCl pH 7.5 (at 20°C), 10 mM DTT), mixed at a molar ratio of 1.2:1.2:1:1, respectively, and dialysed three times against refolding buffer (2 M NaCl, 20 mM Tris-HCl pH 7.5 (at 4°C), 1 mM EDTA, 5 mM β-mercaptoethanol) at 4°C. The formed octamer was purified with a HiLoad 16/600 Superdex 200 pg column (Cytiva). Fractions containing the histones in a equimolar ratio, analysed by SDS-PAGE and Coomassie staining, were pooled, concentrated with a 10,000 MWCO Amicon Ultra Centrifugal Filter (Merck), quantified by 280 nm absorption, and stored at 4°C for use.

### Nucleosome reconstitution

Nucleosome was reconstituted from histone octamer and DNA template at 1.1:1 molar ratio by dialysis from a high salt buffer (“RB high”; (*55*); 1 mM EDTA, 2 M NaCl, 2 mM DTT in 20 mM HEPES, pH 7.5) to a low salt buffer (“RB low”; 1 mM EDTA, 30 mM NaCl, 2 mM DTT in 20 mM HEPES, pH 7.5) over 24 hours (*17*, *55*). Successful reconstitution of the nucleosome was validated by electrophoresis in an 3% agarose gel.

### Cryo-EM sample preparation & data acquisition

Nucleosome-ELF2 samples were prepared by incubating the reconstituted nucleosomes (at 200 nM concentration) with ELF2 protein (1:20 molar ratio) at 25°C for 25 min. Subsequently, 3.2 µl samples of the solution were applied onto EM grids (Quantifoil, R 1.2/1.3 Cu 300 mesh) that had been glow discharged with PELCO easiGlow for 60 seconds in Vitrobot Mark IV chamber at 4°C and 90% humidity. The grid was then blotted for 3 seconds with -5 blotting force and vitrified by plunging into liquid ethane.

Single particle data was collected on Titan Krios G2 microscope (FEI) equipped with a K3 Summit direct electron detector (Gatan). Data was collected using EPU software (ThermoFisher), at nominal magnification of 130,000 x, with a pixel size of 0.652 Å, defocus range from -2.4 to -0.8 at 0.2 intervals, energy filter slit set to 20 eV, and total electron dose of 51.09 electrons/Å^2^ distributed across 44 movie frames.

### Data processing and analysis

Contrast-transfer functions for micrographs were estimated using Warp (*57*), and particles were picked using the RELION 4.0 template picking tool using 2D references generated from a nucleosome map (*58*, *59*). To speed up the initial particle cleanup, the picked particles were extracted and binned to a pixel size of 2.608 Å, and cleaned up using cryoSPARC (*60*) with several rounds of 2D classification, ab-initio reconstruction and heterogeneous refinement. Particles were then imported back into RELION and further cleaned up to include only classes containing a nucleosome by 3D classifications and 3D refinements. A local mask was then made on the trans-DNA at superhelical location +4 to +7 and used for partial signal subtraction, and a 3D classification performed locally with the subtracted particles. The classes with trans-DNA and the class without trans-DNA were selected, and particles re-extracted with pixel size of 1.304 Å.

The region containing ELF2 density was classified after partial signal subtraction and classes with two ELF2 bound to DNA were selected. The selected particles were subsequently refined locally, followed by further rounds of 3D classifications. For an overview of data processing workflow, see **Fig. S2**, for Fourier Shell Correlation (FSC) curves of refined maps, see **Fig. S3**.

### Model fitting and validation

A nucleosome model (PDB 8YBJ (*61*) was fitted into the ELF2-nucleosome cryo-EM map in ChimeraX (*62*) and trimmed and mutated where appropriate with Coot (*63*). An Alphafold3 prediction (*64*) of two ELF2 DBDs bound to the double motif was used for initial rigid body fitting into a map focused on the ELF2 binding sites, with fitting and conservative adjustments made with Coot according to the ELF1 crystal structure described in PDB 8BZM (*65*, *66*). To account for the unwrapped DNA, an ideal B-form DNA double helix was created in Coot. The model was assessed using Phenix (*67*). Solvation free energy and interface area between histones and DNA was calculated using PISA (*68*).

### Electrophoretic mobility shift assay

All steps of EMSA were performed on ice, with running buffer pre-chilled to 4°C. EMSA was performed on an 8% Tris-Glycine gel (Thermo Fisher). 1X Tris-Glycine Native Running Buffer (Thermo Fisher) was used as a running buffer. The gel was pre-rinsed with Running Buffer before assembly of the gel tank. Tris-Glycine Native Sample Buffer (Thermo Fisher) was added to ELF2-nucleosome reaction and mixed by pipetting. Samples were then loaded onto gel and ran on ice at 125V for 2 h. The gel was then stained in SYBR-gold (Thermo Fisher) and visualized using Bio-Rad Imager.

### Restriction enzyme digestion assay

Nucleosomes were reconstituted with modified DNA ligand containing an EcoRV site. Reconstituted nucleosome (containing 0.5 µg of DNA) was then incubated with ELF2 at 1:20 molar ratio at 25°C for 10 minutes, followed by addition of 1.5 U of EcoRV (NEB), and further incubation of the mixture for 15 min at 37°C. To stop the reaction, Proteinase K and SDS were added to 0.2 mg/ml and 0.5% final concentrations, respectively, followed by incubation at 55°C for 10min. The DNA fragments were then analyzed by electrophoresis at a 3% agarose gel.

### Genome mapping of motifs

TF motifs (ETS dual motif, ELF2 single motif , YY1 motif) were obtained from (*8*, *35*, *69*), and their positioning on human genome (hg38) was determined using MOODS (*24*). The relative positions of motifs relative to TSS was determined using HOMER (*70*). Positioning of motifs against TSS was plotted with python Plotly with a 10 bp sliding window.

### CRISPR/Cas9-mediated genome editing

The donor plasmid used for knock-ins contained a GFP-P2A-BSD cassette flanked by two 800 bp homology arms (HAs). These cassettes were PCR-amplified and inserted into the pCMV vector using the HiFi Assembly method (NEB). The ELF2 locus was edited by introducing a CRISPR–Cas9-mediated double-strand break (DSB) and a locus-specific HDR template. Guide RNA sequences were designed using CRISPick (https://portals.broadinstitute.org/gppx/crispick/public), with a preference for protospacers located closest to the stop codon of ELF2. sgRNAs and HDR templates used in this work are provided in **Table S3, S4**.

### Cell lines and transfections, and cell sorting

Human immortalized myelogenous leukemia cells (K562) were kindly provided by Dr. Iva A. Tchasovnikarova (Department of Biochemistry, Gurdon Institute, Cambridge, UK). Cells were cultured in RPMI-1640 (Sigma-Aldrich, catalog no. R8758) supplemented with 10% fetal bovine serum (FBS), 2 nM L-glutamine, and 100 U/mL penicillin-streptomycin, per vendor guidelines. All cell lines were maintained at 37°C with 5% CO₂. Cell authentication was confirmed by suppliers, and lines were tested to be free of mycoplasma contamination.

On the day of transfection, K562 cells were counted using a Countess Automated Cell Counter (Invitrogen), and diluted to 5 × 10⁵ cells/mL in RPMI-1640. 1 mL of diluted cells, supplemented with 0.5 µM NU7441 (Selleck Chemicals) and 10 µM ART558 (MedChem Express), was added to each well of a 12-well plate. For transfection, the following components were combined in 100 µL Opti-MEM (Thermo Fisher Scientific) per well: 750 ng of Cas9 plasmid, 250 ng of sgRNA plasmid, and 1 µg of donor plasmid. 2 µL of PLUS™ Reagent (Thermo Fisher Scientific) was added to the DNA mixture and incubated for 15 minutes. Subsequently, 5 µL of Lipofectamine® LTX was diluted into the mixture, gently mixed, and incubated at room temperature for 25 minutes to form DNA-Lipofectamine complexes. These complexes were then added dropwise to the cells.

Seventy-two hours post-transfection, cells were washed with PBS (Thermo Fisher Scientific) and resuspended in fresh RPMI-1640 containing 10 µg/mL blasticidin (Thermo Fisher Scientific). The medium was replaced every 3–4 days with fresh medium containing blasticidin. After 2 weeks of selection, surviving cells were cultured in fresh medium for two passages to allow recovery. The cells were then washed twice, resuspended in sorting buffer (PBS supplemented with 2% FBS and 1 mM EDTA), and sorted into single cell clones using a BD FACSAria™ III cell sorter (BD Biosciences). The K562 ELF2-GFP knock-in cell lines were genotyped by PCR (primer design see **Table S5**).

### ChIP-seq

For ChIP-seq, we used the two K562 ELF2-GFP knock-in cell lines. ChIP-seq was performed as described previously (*71*, *72*). 8 x 10^6^ cells were harvested and washed with cold PBS and centrifuged at 300 x g for 3 min. The cells were then crosslinked with 10 mL of 1% formaldehyde in RPMI1640 for 10 min, with gentle agitation. The crosslinking was quenched with 1mL 1.25M glycine for 5 min with gentle agitation, and subsequently washed twice with ice cold PBS. the cells were lysed with 0.5 mL lysis buffer (100 mM NaCl, 10mM EDTA, 0.25% Triton X-100 in 10 mM Tris-Cl, pH 8) on ice for 45 min, followed by 0.8mL 1% SDS lysis buffer (150 mM NaCl, 1% SDS, 2 mM EDTA, 1% Triton X-100 and 0.1% sodium deoxycholate in 50mM HEPES, pH 7.5) on ice for 45min and 0.1% SDS lysis buffer (150 mM NaCl, 0.1% SDS, 2 mM EDTA, 1% Triton X-100 and 0.1% sodium deoxycholate in 50mM HEPES, pH7.5) on ice for 45 min twice. Lysed cells were resuspended in 0.2mL 0.1% SDS lysis buffer and sonicated using Bioruptor Diagenode for 9 cycles. 2.5 µL anti-GFP antibody (ab290, abcam) was crosslinked to 50 µL protein G Dynabeads (10003D, Thermofisher) for 3hr at room temperature. 20 µL chromatin was taken as input. Before immunoprecipitation, the chromatin was cleared for 1.5 h at 4°C with 50 µL protein G Dynabeads. Immunoprecipitation was then performed by incubating cleared chromatin in crosslinked antibody-bead in ChIP buffer (10mM EDTA, 100mM NaCl, 1% Triton X-100 and 0.1% sodium deoxycholate in 50mM Tris-Cl, pH 7.5) at 4°C for 18 h. The beads were then washed twice with ChIP buffer, twice with high salt buffer (10 mM EDTA, 500 mM NaCl, 1% Triton X-100, 0.1% sodium deoxycholate in 50 mM Tris-Cl, pH 7.5), twice with Tris/LiCl buffer (0.25M LiCl_2_, 0.5% NP-40, 0.5% sodium deoxycholate and 1 mM EDTA in 10 mM Tris-Cl pH 8), twice with TE buffer (20 mM EDTA in 100 mM Tris-Cl, pH8) at 4°C. DNA was eluted twice in 50 µL elution buffer (10 mM EDTA, 5 mM DTT, 1% SDS in 10 mM Tris-Cl, pH8) at 65°C for 15 min. Sequencing library was prepared using NEBNext® Multiplex Oligos for Illumina® (E7600S, NEB) and NEBNext® Ultra™ II DNA Library Prep Kit for Illumina® (E7645S, NEB). The paired end sequencing (100bpPE) was performed using Illumina® NovaSeqX (CRUK Cambridge Institute genomics core facility).

### ChIP-seq data analysis

The raw sequencing data were adapter trimmed with fastp (v 0.23.4, (*73*)) and quality control was performed with FastQC (v 0.12.1, https://www.bioinformatics.babraham.ac.uk/projects/fastqc/). Trimmed reads were mapped to human genome hg38 using bowtie2 (v 2.5.1, (*74*)). Bam files were created using samtools (*75*). Blacklist regions were removed using intersectBed. Peaks were then called with macs2 (v 2.2.9.1, (*76*)) and visualised using IGV (*77*).

For BPNet analysis, high-confidence binding regions were identified using the IDR tool to determine consensus peaks and summits (*78*). Summits were extended by ±500 bp to generate 1 kb sequences, which were used as training input for the BPNet model (*31*). Regions without signal were included as true negatives to augment the input dataset. The BPNet model architecture (tensorflow version: 1.15.5) consisted of 64 filters and 9 convolutional layers. The model was trained using the Adam optimizer with a learning rate of 0.004 to predict base-resolution binding profiles. Regions from chromosome 11 were used as the test set, while regions from chromosomes 1 and 2 were used for validation during hyperparameter tuning. ChIP control samples were incorporated to account for biases and background noise. The DeepLIFT tool (v 0.1) was used to compute the contribution scores (*79*) which were then analyzed with the TF-MoDISco tool (v 0.5.3.0) to identify significant seqlets and motif discovery (*80*) .

De novo motif discovery was performed using HOMER (v 5.1, (*70*)). TSS position was identified using HOMER annotatePeaks.pl. The peak-TSS position heatmap (**Fig. 4b**) is plotted using deeptools (*81*). Gene Ontology analysis was performed using Metascape (*82*). The gene expression level analysis was performed using K562 gene expression level data from DEPMAP (https://doi.org/10.25452/figshare.plus.24667905.v2), the genes were grouped into 3 groups: no ETS-HT2 double motif near the TSS of the gene (no motif, grey), have ETS-HT2 double motif within 20 bp of the TSS and in the orientation that it would make TSS accessible (correct orientation, red) and everything else (yellow). The ETS-HT2 double motif positions used in the gene expression analysis were the motifs found within ChIP-seq peaks, and the plot was plotted using Python. For the methylation analysis, the whole genome bisulfite sequencing methylation data of K562 was taken from GEO (accession GSE86747)(*36*), motif methylation was analysed using bedtools (*83*) where all ETS-HT2 motifs and ELF2 single motifs mapped on both ELF2-ChIP-seq and methylation data are taken and sorted by methylation and ChIP-occupancy respectively. For the double motifs, only no methylation on either sites is counted as non-methylated and methylation detected on both sites are counted as methylated. The percentage of motifs found in ChIP-seq peaks are plotted as an cumulative percentage of all motifs include and above the score against motif score from MOODS. The expression levels of the genes were plotted as scatter plot and box plot using python and p-value was determined with 2-sample, independent Student’s t-test.

### mC-SELEX experiment and data analysis

The full-length recombinant human ELF2 protein was expressed and purified as described in (*35*). In brief, a Gateway recipient vector containing an N-terminal Thioredoxin- 6× His tag and a 3× FLAG tag was used for bacterial expression, with pETG20A plasmid serving as backbone. The recombinant protein was produced and purified from E. coli and analysed by SDS-PAGE electrophoresis (Invitrogen, EP09606), followed by PageBlue staining (Thermo Scientific, 24620) to confirm the expression. The purified protein was supplemented with 50% glycerol and stored at –20°C.

The Methylcytosine Systematic Evolution of Ligands by Exponential Enrichment (mC- SELEX) was performed at four different 5-methylcytosine (5-mC) levels. The mC-SELEX experiment followed previous methods (*35*), with modifications to sequencing adapters and inclusion of 5-mC. In brief, DNA ligands were designed with a 40 bp random sequence flanked by an 8 bp barcode and Illumina TruSeq adapters, and the barcode used has been described in a previous study (*8*). DNA ligands were amplified by PCR before each mC-SELEX cycle, during which 5-mC was introduced by mixing 5-methyl-dCTP (New England Biolabs, N0356S) with regular dCTP (Thermo Fisher, R0193) in certain proportions. These included four setups: 5- methyl-dCTP:dCTP = 0:3, 1:2, 2:1, 3:0, corresponding to 0% mC, 33% mC, 67% mC, and 100% mC, respectively. The DNA ligands were then incubated with the protein in low-salt incubation buffer (4% glycerol, 1 mM DTT, 100 μM EGTA, 8.33 μg/mL Poly(dI-dC), 1 mM MgCl_2_, 3 μM ZnSO_4_, 75 mM NaCl in 20 mM Tris-Cl, pH 7.5) to allow protein-DNA binding reactions followed by pull down of the proteins with bound DNA with nickel affinity beads (Cytiva, 28967390). The DNA ligands were then eluted and purified using AMPure XP Reagent (Beckman Coulter, A63881), and sequenced using DNBSEQ-G400 (BGI Genomics).

To characterise the binding specificity of transcription factors, motif logos were generated from PWM based on the enriched sequences from mC-SELEX. The binding sites were identified using the Autoseed algorithm (*84*), and each PWM was retrieved by aligning the most enriched motifs from the final cycle of mC-SELEX. These models were then validated by comparing to previously characterised motifs in databases, JASPAR (*85*) and the Human Transcription Factors (*4*).

### Competitive Genome Editing assay

HAP1 (#C631) cell line was obtained from Horizon Discovery and maintained in low-density cultures in Iscove’s Modified Dulbecco’s Medium (IMDM) according to the vendor’s guidelines. Each genomic locus was edited via competitive genome editing (CGE) assay as described in (*39*). In brief, 200,000 early passage HAP1 cells per well in 12-well plate were transfected with RNP complexes containing sgRNA (250 ng, Integrated DNA technologies, **Table S3, S4, S6**) and S.p. HiFi Cas9 protein (1000 ng, Integrated DNA technologies) with single stranded HDR templates at final concentration of 3 nM using CRISPRMAX (Thermo Fisher Scientific # CMAX00008) following manufacturer’s instructions. Analysis of fitness effects were performed as described in (*39*), where fold change in cell fitness was calculated as log2[day 7 read count (mutated/original)/day 2 read count (mutated/original)].

## Data availability

The Cryo-EM maps and final model have been deposited in the Electron Microscopy Data Bank (EMD-52852) and the Protein Data Bank (9IGJ). The sequencing data will be deposited to European Nucleotide Archive (accession: PRJEB85947).

## ACKNOWLEDGMENTS

We thank Kenneth Jones for technical assistance, and Prof. Ben Luisi for critical review of the manuscript. Part of this work was facilitated by the Protein Science Facility at Karolinska Institutet/SciLifeLab (http://ki.se/psf) and we would like to thank Dr. Henry Ampah-Korsah and Dr. Emilia Strandback for the assistance. We thank Mr Lee R. Cooper, Dr Steve W. Hardwick and Dr Dimitri Y. Chirgadze for their assistance using the Cryo-EM Facility at the Department of Biochemistry, University of Cambridge. We thank the flow cytometry facility from the School of Biological Sciences, University of Cambridge for their support and assistance in cell sorting. We thank the Cancer Research UK Cambridge Institute Genomics Core facility for their assistance in DNA sequencing. This work was supported by the Max Planck Society, Cancer Research UK (Grant number RG99643) and UK Research and Innovation (Biotechnology and Biological Sciences Research Council G107673 and Medical Research Council G105296).

## Notes

### Competing Interest Statement

The authors have declared no competing interest.

### Summary of Updates

updated title; updated authors list and affiliations; added new in vivo results; Figures updated with addition of in vivo results; supplementary files updated

