## Supplementary Figures for "The pioneer transcription factor ELF2 remodels the nucleosome near transcription start sites"

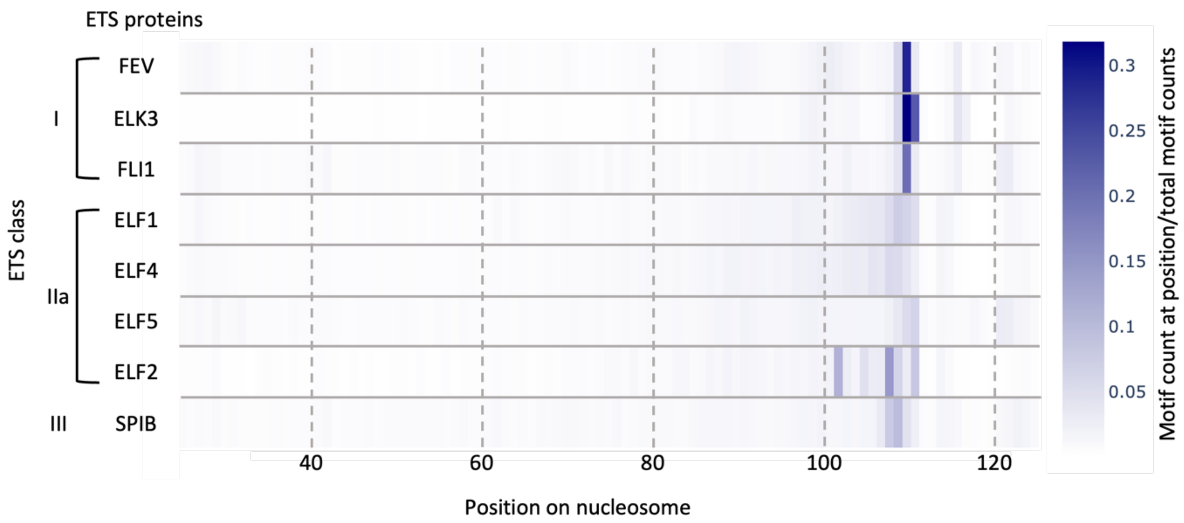


**Fig. S1: ETS family protein motif localisation on nucleosome.**

Heatmap of preferred double motif binding location of ETS family TFs on + strand of nucleosomal DNA. Scale shows fraction of binding motif matches relative to all matches for that factor within the nucleosome. Data from (Zhu et al., 2018).

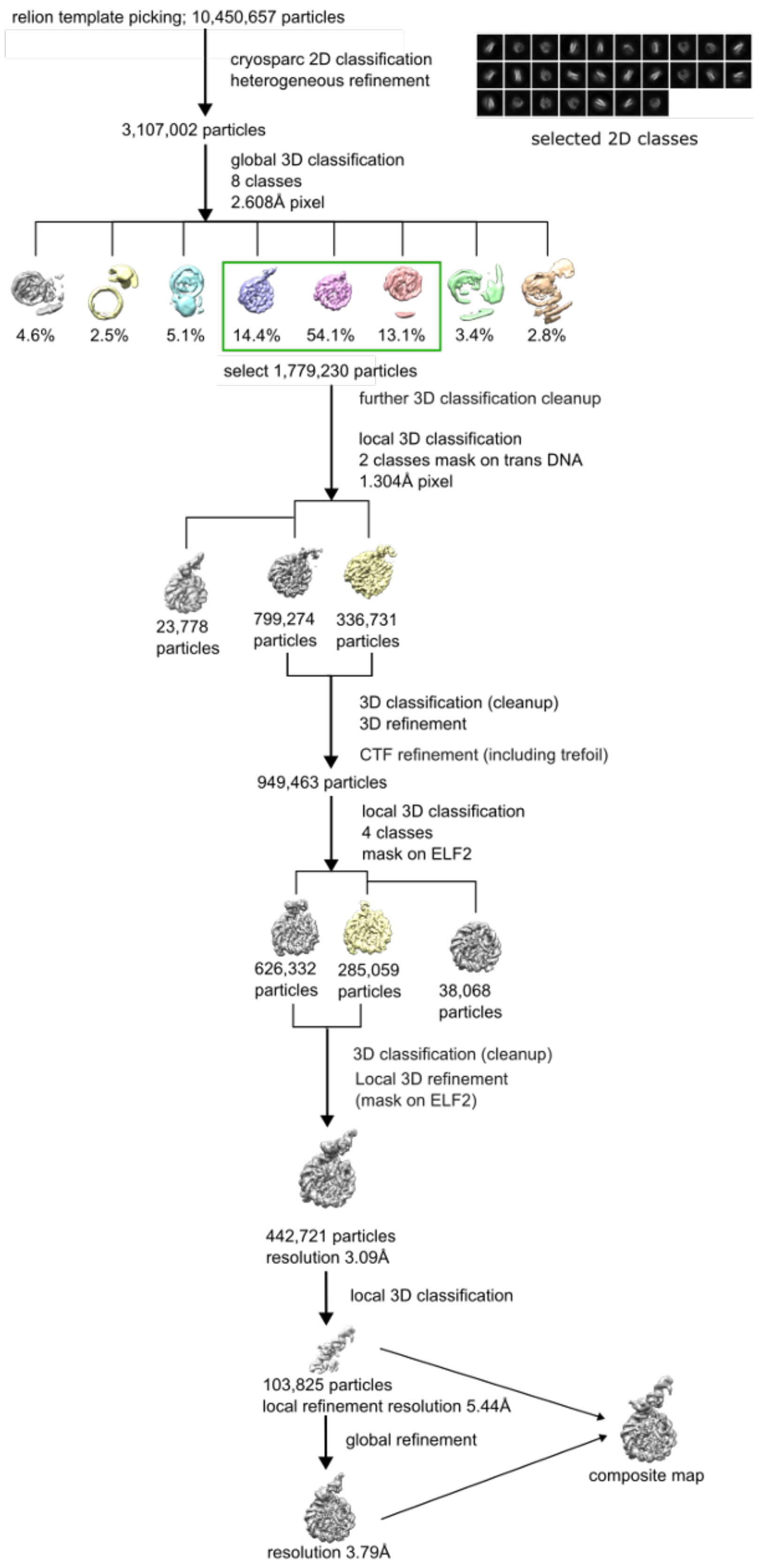


**Fig. S2: Flowchart of the cryo-EM data analysis workflow.**

Figure shows data processing workflow for solving the ELF2-nucleosome structure.

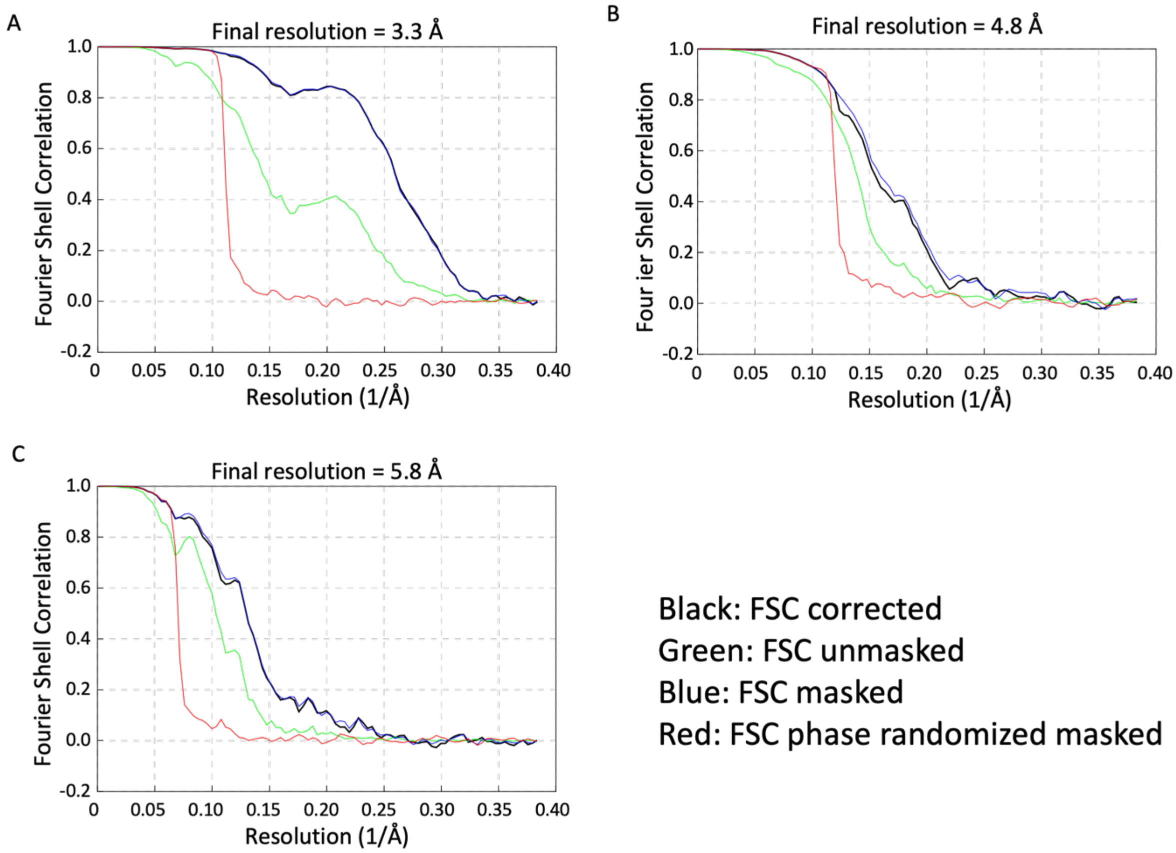


**Fig. S3: FSC curves.**

**A**: FSC for global refinement in **Fig. 2a** and **Fig. S4c**.

**B**: FSC for local refinement in **Fig. 2a** and **Fig. S4a**.

**C**: FSC for map in **Fig. S4d**.

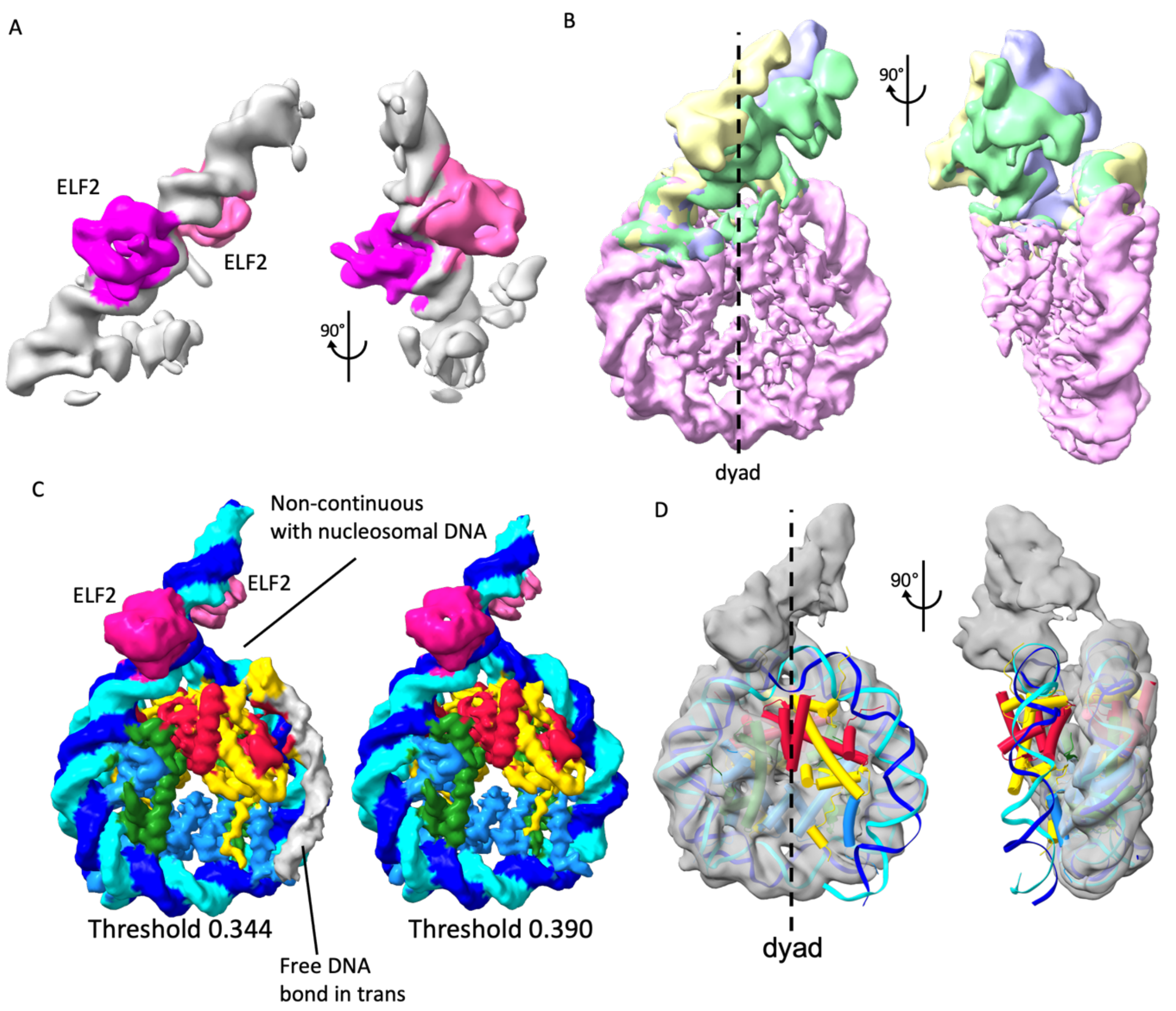


**Fig. S4: ELF2-nucleosome map and unbound nucleosome map.**

A: Front and side view of locally refined map (103,825 particles) around ELF2-bound DNA. Two ELF2 densities bound on DNA are highlighted.

B: Front and side view of three local 3D classifications around the ELF2 binding region are shown, overlayed onto a canonical nucleosome map, showing the flexibility of unbound DNA.

C: The left panel shows the refinement map at a threshold of 0.344, where the trans-DNA can be observed in the cryo-EM map (grey). The right panel is the same volume shown with a threshold of 0.390, where the trans-DNA is no longer visible.

D: Structure of ELF2-bound hexasome. Cryo-EM map with canonical nucleosome (6FQ5) fitted is shown.

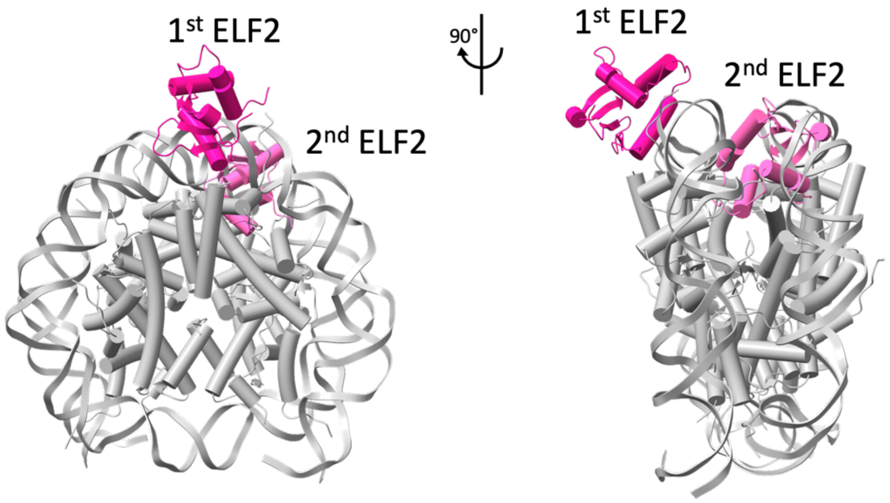


**Fig. S5: Model of ELF2 binding the ELF2-HT2 motif in the context of a canonical nucleosome (6FQ5).**

A model of two ELF2 proteins bound to the ELF2-HT2 motif on a nucleosome where the DNA position is similar to a canonical nucleosome is shown. Note that the second ELF2 binding site is not accessible as that ELF2 would clash with the nucleosome when DNA is not unwrapped.

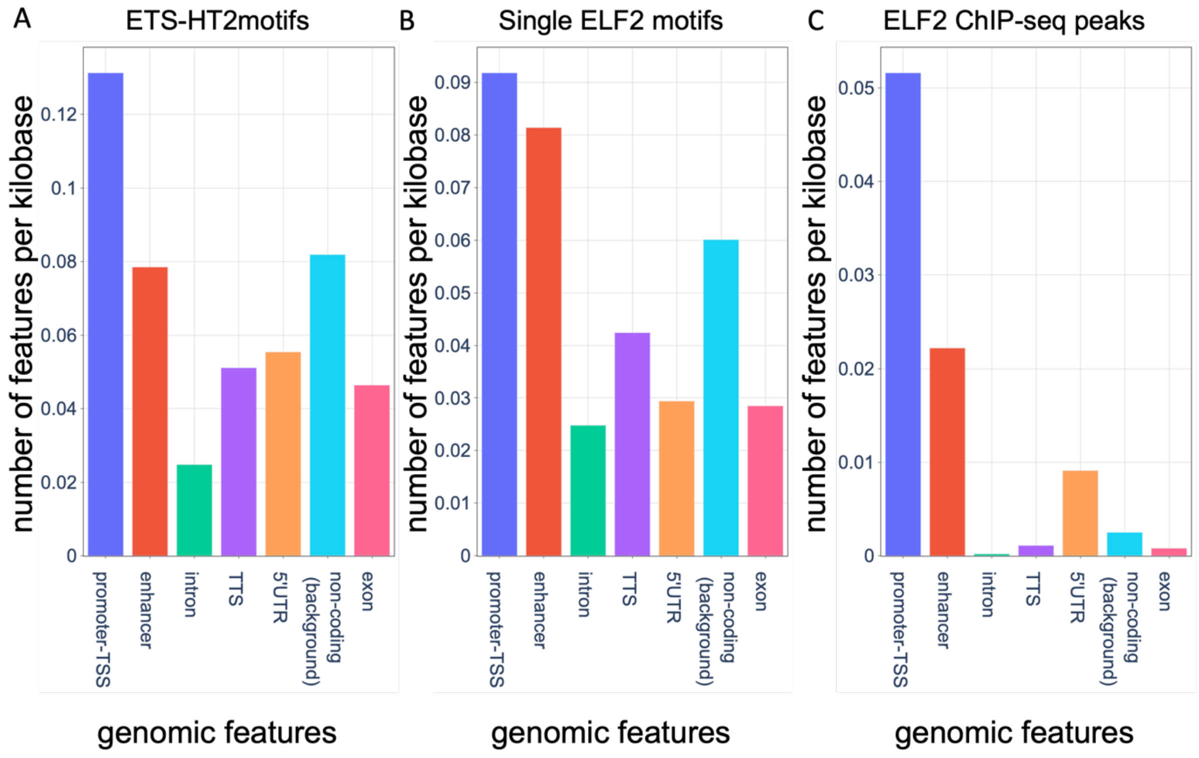


**Fig. S6: Bar plots of genomic features.**

A: Bar plot showing the occurrence of ETS-HT2 motifs at different genomic features per kilobase. 300,000 motifs were selected from hg38 genome using moods, and the overlap with genomic features were counted using HOMER, the enhancer annotation was obtained from (Lidschreiber et al., 2021).

B: bar plot showing the occurrence of ELF2 single motifs at different genomic features per kilo-basepair.

C: bar plot showing the occurrence of ELF2-GFP ChIP-seq peaks at different genomic features per kilobase.

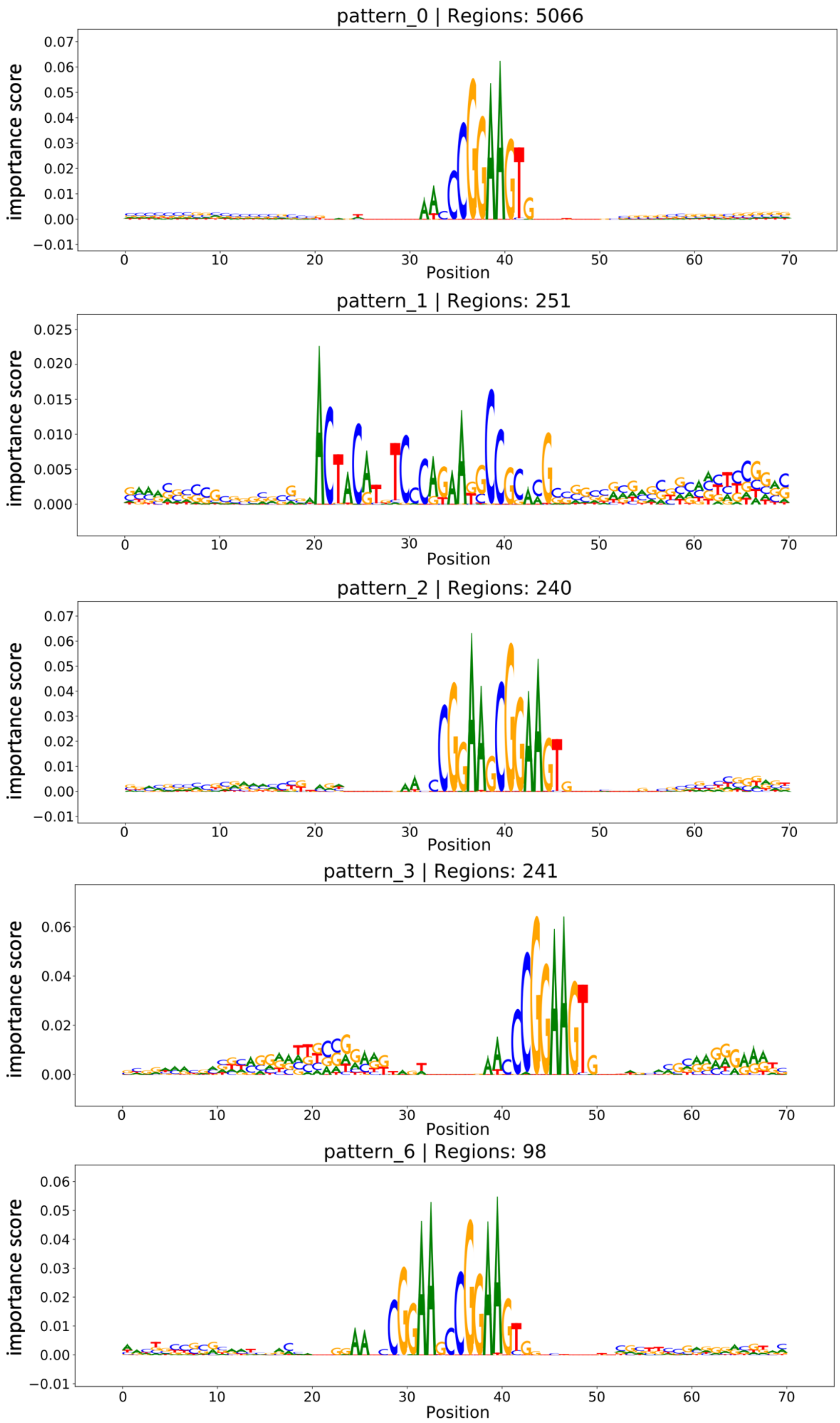


**Fig. S7: Motif logos from ChIP-seq generated using BPNet.**

Sequence logos representing aggregated contribution scores for the identified ELF2-associated motifs from BPNet. The y-axis shows the average contribution scores of each nucleotide to the model's prediction across all seqlets while the x-axis shows the nucleotide position.

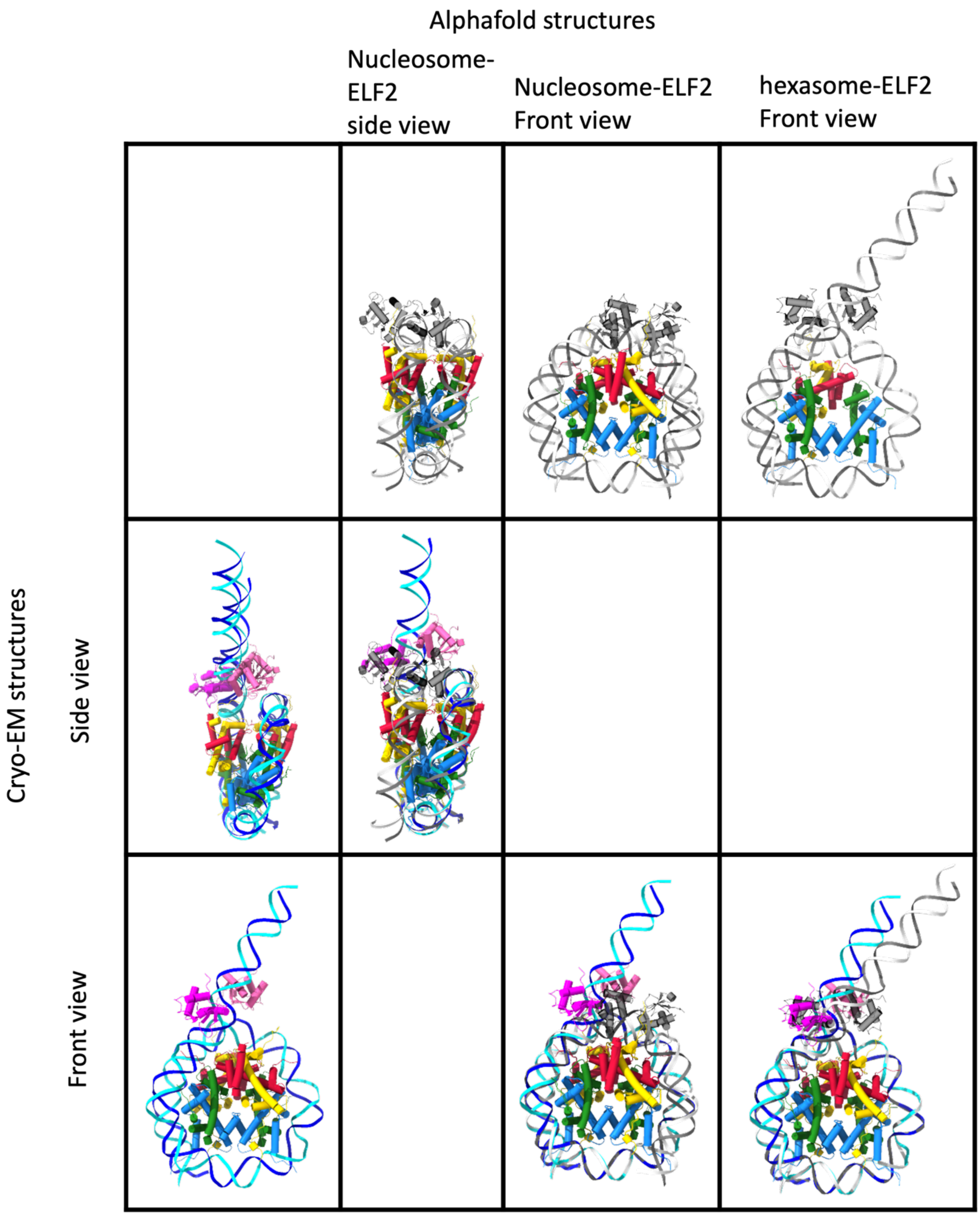


**Fig. S8: Alphafold prediction of ELF2-nucleosome interactions.**

The figure shows Alphafold predictions using Alphafold 3. In the first row, the figures show the Alphafold prediction of the side and front view of nucleosome interaction with two ELF2 proteins and the front view of hexasome interaction with two ELF2 proteins. In the first column the figure shows the side and front view of the ELF2-nucleosome model from Cryo-

EM. The second and third row of the figure shows the overlay between the Alphafold predicted structure and Cryo-EM structure.
