## Supplementary Tables for "The pioneer transcription factor ELF2 remodels the nucleosome near transcription start sites"

Table S1: Cryo-EM data collection, model building and refinement statistics

| | |
|---|---|
| **Data Collection** | |
| Magnification (×) | 130,000 |
| Electron fluence (e−/Å2) | 51.09 |
| Defocus range (µm) | -2.4 to -0.8 |
| Pixel size (Å) | 0.652 |
| Initial particles | 10,450,657 |
| nucleosome-like particles after initial cleanup | 1,179,230 |
| Final particles | 442,721 |
| Average resolution (Å) (FSC threshold 0.143) | 3.29 |
| **Model composition** | |
| Atoms | 12532 |
| Protein residues | 920 |
| nucleotides | 250 |
| **Refinement** | |
| Map CC around atoms | 0.74 |
| Map sharpening B factor (Å2) | -137 |
| R.M.S. deviations | |
| Bond lengths (Å) | 0.007 |
| Bond angles (°) | 1.106 |
| **Validation** | |
| MolProbity score | 1.01 |
| Clash score | 2.31 |
| Poor rotamers (%) | 0.77 |
| Ramachandran statistics | |
| Favoured (%) | 98.56 |
| Outlier (%) | 0.44 |

Table S2: H2A and H2B solvation free energy and interface area interacting with DNA. The data is obtained using PISA.

| | ΔG (kcal/mol) | | interface area (Å$^2$) | |
|---|---|---|---|---|
| | ELF2-nuc | nuc | ELF2-nuc | nuc |
| H2A interaction with + strand | -1.7 | -3.5 | 79.9 | 531.5 |
| H2A interaction with - strand | - | -5.8 | - | 504.9 |
| H2B interaction with + strand | -3.1 | -5 | 161.2 | 440.7 |
| H2B interaction with - strand | -0.2 | -6.5 | 38.5 | 400.1 |

Table S3: sgRNA design

| | |
|---|---|
| | sgRNA Sequence |
| ELF2-GFP-knockin | ATAAAATAGCAGCTCCACCA |
| | crRNA sequence |
| CGE-negative-control | TGTTGGTGAAGCTAACGTTG |
| CGE-RPL23 | CCCCGCCCACCCTCCTCACG |
| CGE-EIF3K | GCGTCCTCAAGGACGGGAAC |

Table S4: HDR templates

| name | HDR template |
|---|---|
| ELF2-GFP-P2A-BSD | TAGTTTTTAAAATATATCATGGTATGTTTGTGTCAAACATCAGGGATTGCCAGTACC<br>TCTGTAAAATTTGGGAGCACTAATTTTACACATTTTGGATTTGTATTAATCTTTTACT<br>CAGGAGACCTATTTTCTCAGTTGGCATTAAATATAATCAGCTTTTTTCTTTCTAATGT<br>ACTAGGACAGTTCGTGTGGCAATGCAGGTACCTGTTGTAATGACATCATTGGGTCA<br>GAAAATTTCAACTGTGGCAGTTCAGTCAGTTAATGCAGGTGCACCATTAATAACCA<br>GCACTAGTCCAACAACAGCGACCTCTCCAAAGGTAGTCATTCAGACAATCCCTACT<br>GTGATGCCAGCTTCTACTGAAAATGGAGACAAAATCACCATGCAGCCTGCCAAAAT<br>TATTACCATCCCAGCTACACAGCTTGCACAGTGTCAACTGCAGACAAAGTCAAATCT<br>GACTGGATCAGGAAGCATTAACATTGTTGGAACCCCATTGGCTGTGAGAGCACTTA<br>CCCCTGTTTCAATAGCCCATGGTACACCTGTAATGAGACTATCAATGCCTACTCAGC<br>AGGCATCTGGCCAGACTCCTCCTCGAGTTATCAGTGCAGTCATAAAGGGGCCAGA<br>GGTTAAATCGGAAGCAGTGGCAAAAAAGCAAGAACATGATGTGAAAACTTTGCAG<br>CTAGTAGAAGAAAAACCAGCAGATGGAAATAAGACAGTGACCCACGTAGTGGTTG<br>TCAGTGCGCCTTCAGCTATTGCCCTTCCTGTAACTATGAAAACAGAAGGACTAGTG<br>ACATGTGAGAAAAACGGATCCGGCAGCGGCGGGGAATTCATGGTGAGCAAGGGC<br>GAGGAGCTGTTCACCGGGGTGGTGCCCATCCTGGTCGAGCTGGACGGCGACGTAA<br>ACGGCCACAAGTTCAGCGTGTCCGGCGAGGGCGAGGGCGATGCCACCTACGGCA<br>AGCTGACCCTGAAGTTCATCTGCACCACCGGCAAGCTGCCCGTGCCCTGGCCCACC<br>CTCGTGACCACCCTGACCTACGGCGTGCAGTGCTTCAGCCGCTACCCCGACCACAT<br>GAAGCAGCACGACTTCTTCAAGTCCGCCATGCCCGAAGGCTACGTCCAGGAGCGC<br>ACCATCTTCTTCAAGGACGACGGCAACTACAAGACCCGCGCCGAGGTGAAGTTCG<br>AGGGCGACACCCTGGTGAACCGCATCGAGCTGAAGGGCATCGACTTCAAGGAGG<br>ACGGCAACATCCTGGGGCACAAGCTGGAGTACAACTACAACAGCCACAACGTCTA<br>TATCATGGCCGACAAGCAGAAGAACGGCATCAAGGTGAACTTCAAGATCCGCCAC<br>AACATCGAGGACGGCAGCGTGCAGCTCGCCGACCACTACCAGCAGAACACCCCCA<br>TCGGCGACGGCCCCGTGCTGCTGCCCGACAACCACTACCTGAGCACCCAGTCCGCC<br>CTGAGCAAAGACCCCAACGAGAAGCGCGATCACATGGTCCTGCTGGAGTTCGTGA<br>CCGCCGCCGGGATCACTCTCGGCATGGACGAGCTGTACAAGGGATCCGGCGCAAC<br>AAACTTCTCTCTGCTGAAACAAGCCGGAGATGTCGAAGAGAATCCTGGACCGATG<br>GCCAAGCCTTTGTCTCAAGAAGAATCCACCCTCATTGAAAGAGCAACGGCTACAAT<br>CAACAGCATCCCCATCTCTGAAGACTACAGCGTCGCCAGCGCAGCTCTCTCTAGCG<br>ACGGCCGCATCTTCACTGGTGTCAATGTATATCATTTTACTGGGGGACCTTGTGCA<br>GAACTCGTGGTGCTGGGCACTGCTGCTGCTGCGGCAGCTGGCAACCTGACTTGTAT<br>CGTCGCGATCGGAAATGAGAACAGGGGCATCTTGAGCCCCTGCGGACGGTGCCG<br>ACAGGTGCTTCTCGATCTGCATCCTGGGATCAAAGCCATAGTGAAGGACAGTGAT<br>GGACAGCCGACGGCAGTTGGGATTCGTGAATTGCTGCCCTCTGGTTATGTGTGGG<br>AGGGCTAACCATGGACTTCAGGCTGTTAGTGGCAGTACTGACATAAACATTTGCAA<br>GGGAAGTCATCAAGAAAAGTCAAAGAAGACTTTAAAACATTTTTAATGCATATACA<br>AAAACAATCAGACTTACTGGAAATAAATTACCTATCCCATGTTTCAGTGGGAAATG<br>AACTACATATTGAGATGCTGACAGAAAACTGCCTCTTACAGTAGGAAACAACTGAA<br>CCCATCAATAAGAAAAAGGATCGAAAGGGACCAAGCAGCTCACTACGATATCAAG<br>TTACACTAAGACTTGGAACACTAACATTCTGTAAGAGGTTATATAGTTTTCAGTGG<br>GAGGGGTTGGGATGGGTAATCTCATTGTTACATATAGCAATTTTTGATGCATTTTAT<br>ATGCATACCAGCAATTATTACTGTGTTCGCACAGTTCTCACTTAACTGGTGCTATGT |

| | |
|---|---|
| | GAAGACTCTGCTAATATAGGTATTTTAGAATGTGAATTGAAGAATGGATCCCAAAA ACTTCAGAAAGAGGATAGCAAAAAAAGATCTAGTGCGATTTTATATATATATATAT ATATATATACATACATATATATATATCATATAGCTTAAGCTGATTTAAAACAAAGGC CTTAGACTAATTTTCGATTTTCTTTCTTGAAATAAGCTAATGGCTTGTTTGTGTAAAG CTTTTTTATTAAAAGAAAAATTTTAAAAATCTTGTACCTAGCACAGTATTGTTATAG AATATACATGTAACATTTTATATGGTAGTTTAAGTCTGTCAGTTTCTTAATTGTGGA CAAATTAACAGTTGGCTC |
| Negative control-Original | TACGGCTGCACCGAGTCGTAGTCGAGGTCATAGTTCCTGTTNGTNAAGCTAACNTT NAGGGGCATCGTCGC GGGAGGCTGCTGGAGCGGGGCACACAAAG |
| Negative control-Mutated | TACGGCTGCACCGAGTCGTAGTCGAGGTCATAGTTCCTGTTNGTNAAACTTACNTT NAGGGGCATCGTCGC GGGAGGCTGCTGGAGCGGGGCACACAAAG |
| RPL23-Original | TAATAAGGCAGCGCCCAGAGGCGGAAGAGGCCGGTTTTTGCT (N4:08760808)(N4) (N3:08087608)(N3)(N4) CACGTG (N1:76080808)(N3)(N3)(N1)(N3) GGTGGGCGGGGCGTTAAAGTTCATATCCCAGTGTCCTTTGAA |
| RPL23-Mutated | TAATAAGGCAGCGCCCAGAGGCGGAAGAGGCCGGTTTTTGCT (N4:08760808)(N4) (N3:08087608)(N3)(N4) TAAATA (N1:76080808)(N3)(N3)(N1)(N3) GGTGGGCGGGGCGTTAAAGTTCATATCCCAGTGTCCTTTGAA |
| EIF3K-Original | GGCTAACACTGACCCGGCACGGCGTCCTCAA(N3:08087608)(N3)(N1:76080808) (N4:08760808)(N3)GGAACAGGAAGAGGTGGTGGAAACGGAA(N1)(N4)(N3)(N1 )(N4)GTCGTTGTTGTCGCTCCAGCTCCTTGGGTGC |
| EIF3K-Mutated | GGCTAACACTGACCCGGCACGGCGTCCTCAA(N3)(N3)(N1)(N4)(N3) GCAACAGCAAGAGGTGGTGCAAACGCAA (N1)(N4)(N3)(N1)(N4)GTCGTTGTTGTCGCTCCAGCTCCTTGGGTGC |

Table S5: primer design for K562 ELF2-GFP knock-in cell line genotyping

| primer | sequence |
|---|---|
| 1_fw | GAACCCCATTGGCTGTGAGA |
| 1_rev | TGCTTGGTCCCTTTCGATCC |
| 2_fw | TGCAGTCATAAAGGGGCCAG |
| 2_rev | CCAACCCCTCCCACTGAAAA |
| 3_fw | CCATCCCAGCTACACAGCTT |
| 3_rev | TCAGTACTGCCACTAACAGCC |

Table S6: Target specific primers for gDNA amplification in CGE

| name | sequence |
|---|---|
| NegCtr-gDNA-FP | /5Biosg/ACACGACGCTCTTCCGATCTCCGCACCAAGACCCCTTTAA |
| NegCtr-gDNA-RP | GACGTGTGCTCTTCCGATCTTCCTCCTCGTCGCAGTAGAA |
| RPL23-gDNA-FP | /5Biosg/ACACGACGCTCTTCCGATCTGCTTCGACATCTTGAACGCC |
| RPL23-gDNA-RP | GACGTGTGCTCTTCCGATCTTGCGCTTCTACCTCAACTCC |
| EIF3K-gDNA-FP | /5Biosg/ACACGACGCTCTTCCGATCTCCTGCGAGGAAGAAGTCAC |
| EIF3K-gDNA-RP | GACGTGTGCTCTTCCGATCTTGGCTCTCATCTGCTCAAAC |
